# Neural signatures of task-related fluctuations in auditory attention change with age

**DOI:** 10.1101/2022.04.12.487991

**Authors:** Björn Herrmann, Burkhard Maess, Molly J. Henry, Jonas Obleser, Ingrid S. Johnsrude

**Author notes:** Correspondence concerning this article should be addressed to Björn Herrmann, Rotman Research Institute, Baycrest, 3560 Bathurst St, North York, Ontario, M6A 2E1, Canada.

## Abstract

Listening in everyday life requires attention to be deployed dynamically – when listening is expected to be difficult and when relevant information is expected to occur – to conserve mental resources. Conserving mental resources may be particularly important for older adults who often experience difficulties understanding speech. In the current study, we use electro- and magnetoencephalography to investigate the neural and behavioral mechanics of attention regulation during listening and the effects that aging has on these. We show that neural alpha oscillatory activity indicates when in time attention is deployed (Experiment 1) and that deployment depends on listening difficulty (Experiment 2). Older adults (54–72 years) also show successful attention regulation but appear to utilize timing information differently compared to younger adults (20–33 years). We further show a notable age-group dissociation in recruited brain regions. In younger adults, superior parietal cortex underlies alpha power during attention regulation, whereas, in older adults, alpha power emerges from more ventro-lateral areas (posterior temporal cortex; Experiment 3). This difference in the sources of alpha activity between age groups only occurred during task performance and was absent during rest (Experiment S1). In sum, our study suggests that older adults employ different neural control strategies compared to younger adults to regulate attention in time under listening challenges.

## Introduction

Speech is often difficult to understand, either because of situational demands (e.g., background speech), or because of age-related changes in sensory systems (Pichora-Fuller, 2003; Mattys et al., 2012; Johnsrude and Rodd, 2016). Degraded speech increases the need to recruit cognitive resources such as attention, but this is mentally costly (Westbrook and Braver, 2015; Hornsby et al., 2016; Shenhav et al., 2017) and makes listening effortful (Pichora-Fuller, 2016; Strauss and Francis, 2017; Peelle, 2018; Herrmann and Johnsrude, 2020). Critically, situational demands vary over time as the intensity and character of background speech change, and not all words are equally important for a listener to follow a conversation. Hence, the effort required to comprehend may also fluctuate. A listener could attend specifically at times when listening is expected to be difficult or when relevant information is expected to occur in order to reduce experienced effort and conserve mental resources. The current work explores the neural and behavioral signatures of attention during listening and the changes associated with aging.

Electro-/magnetoencephalography (EEG/MEG) measuring neural oscillatory activity in the alpha (∼10 Hz) frequency band provides a unique window on cognition (Klimesch, 1999; Klimesch et al., 2007; Jensen and Mazaheri, 2010). High alpha power has been associated with the recruitment of cognitive resources during listening (Adrian, 1944; Foxe et al., 1998; Fu et al., 2001; Palva and Palva, 2011). For example, alpha power increases when individuals listen actively compared to passively (Dimitrijevic et al., 2017; Henry et al., 2017) or listen to degraded, difficult-to-understand speech (Wöstmann et al., 2015). Alpha activity further appears to be sensitive to temporal dynamics of cognitive processing. Alpha power synchronizes with attended words presented in dichotic speech-listening tasks (Wöstmann et al., 2016; Tune et al., 2018; Wöstmann et al., 2021) and temporally aligns with an anticipated visual stimulus (Rohenkohl and Nobre, 2011; Bonnefond and Jensen, 2012; Payne et al., 2013). Here, we focus on how alpha activity changes over time as individuals direct attention to times at which key information is expected to occur during challenging listening.

Older adulthood is associated with reduced cognitive control abilities, including attentional control (Braver and Barch, 2002; Braver, 2012; Zanto and Gazzaley, 2017). Consistent with age-related reductions in cognitive control, behavioral performance and attention-related lateralization of alpha power in visuospatial tasks is reduced in older compared to younger adults (Zanto et al., 2011; Sander et al., 2012; Hong et al., 2015; Leenders et al., 2018; Dahl et al., 2019). In active listening tasks, alpha power typically increases relative to baseline (Wöstmann et al., 2015; Dimitrijevic et al., 2017; Henry et al., 2017), but older adults show weaker power increases (Wöstmann et al., 2015; Rogers et al., 2018) or even suppression of alpha power compared to younger individuals (Henry et al., 2017; Getzmann et al., 2020). Moreover, attention-related lateralization of alpha power in auditory spatial tasks appears to be reduced in older compared to younger adults (Getzmann et al., 2020) and in adults with hearing impairment compared to those without (Bonacci et al., 2019). However, from these works it is unclear whether attending to key points in time during challenging listening is affected by aging.

In the current experiments using EEG (Experiments 1 and 2) and MEG (Experiments 3 and S1), participants listened to 10-s white noise sounds, detecting a gap target that they were informed occurred in the first half of the sound, or in the second half. This paradigm allows us to capture neural activity and behavior as listeners flexibly deploy attention either early or late in a stimulus. We test whether neural alpha power indexes the deployment of attention during listening (Experiment 1) and whether alpha power changes with the degree of expected listening difficulty (Experiment 2). We then investigate whether attending during challenging listening, as indexed by alpha power and behavior, differs between younger and older adults (Experiment 3), and whether any age differences are observed only when a task is performed, or are also present during resting-state alpha recording (Experiment S1).

## General Methods

### Participants

In total, 132 male and female adults of varying age participated in the experiments of the current study. Different participants were recruited for each of the three main experiments and the one supplementary experiment (see Experiment S1 in the Supplemental Materials). Demographic details are described below for each experiment individually. Participants reported no neurological disease or hearing impairment and were naïve to the purpose of the experiment. Participants gave written informed consent prior to the experiment and were paid for their participation. The study was conducted in accordance with the Declaration of Helsinki, the Canadian Tri-Council Policy Statement on Ethical Conduct for Research Involving Humans (TCPS2-2014), and approved by the Nonmedical Research Ethics Board of the University of Western Ontario (protocol ID: 106570).

### Stimulation apparatus

All EEG sessions (Experiments 1 and 2) were carried out in a sound-attenuating booth. Sounds were presented via Sennheiser (HD 25-SP II) headphones and a Steinberg UR22 (Steinberg Media Technologies) external sound card. Stimulation was controlled by a PC (Windows 7) running Psychtoolbox in MATLAB (MathWorks Inc.).

All MEG sessions (Experiments 3 and S1) were carried out in an electromagnetically shielded, sound-attenuating room (Vacuumschmelze, Hanau). Sounds were presented via in-ear phones and the stimulation was controlled by a PC (Windows XP) running Psychtoolbox in MATLAB.

### Acoustic stimulation and procedure

At the beginning of Experiments 1-3, each participant’s hearing threshold was measured for a white noise stimulus using a methods of limits procedure, which we have described in detail in previous work (Herrmann and Johnsrude, 2018). Acoustic stimuli were presented at 50 dB (Experiments 1 and 2) or 45 dB (Experiment 3) above the individual’s hearing threshold (sensation level).

Acoustic stimuli during the main experimental procedures were 10-s white noise sounds that each contained one short gap (Figure 1A). The participants’ task was to press a button as soon as they detected the gap (Henry and Obleser, 2012; Henry et al., 2014, 2016; Henry et al., 2017). The white noise 0.4 s prior to and 0.1 s after a gap was identical for each sound, whereas the remainder of the 10-s white noise differed from trial to trial. This ensured that the probability of detecting a gap would be unaffected by trial-by-trial acoustic variations just prior and subsequent to a gap. Gap duration was titrated for each individual to approximately 70-80% detected gaps in 2-4 training blocks (∼2.5 min each) prior to the main experiment. The noise bursts presented during training each incorporated two gaps presented at random times at least 2 s apart, and participants had to press a response button as soon as they detected a gap.

**Figure 1:**
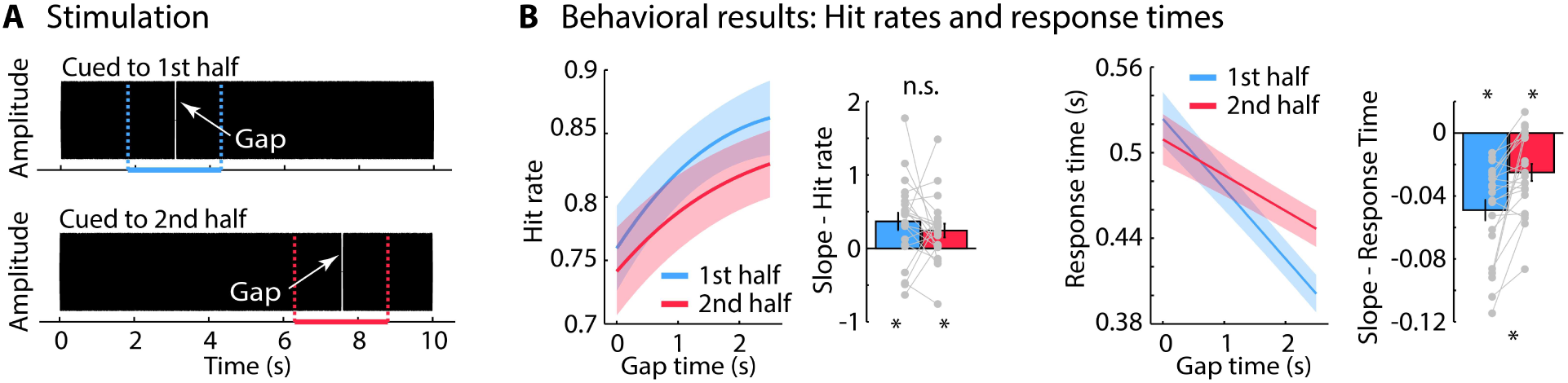
Stimulus representation and behavioral performance for Experiment 1. A: Participants were presented with 10-s white noises that contained one gap. The gap occurred within a 2.5-s window either within the first half (1.8–4.3 s) or within the second half (6.3–8.8 s) of the noise (colored dashed lines). Participants were cued as to whether the gap would occur in the first or the second half of the sound, allowing them to orient their attention in time. Participants did not know exactly when, within the cued half of the sound, the gap would occur. B: Predicted hit rates and response times from logistic and linear fits, respectively, as a function of gap occurrence within the cued 2.5-s window. The x-axis reflects the gap time relative to the onset of the cued time window (1.8 s and 6.3 s for first half and second half, respectively). Slopes from logistic (hit rate) and linear (response time) fits are plotted as bar graphs and individual data points. Error bars and shadings reflect the standard error of the mean. *p ≤ 0.05, n.s. – not significant.

For the main experiment, the gap occurred within a 2.5-s window either within the first half (1.8–4.3 s) or within the second half (6.3–8.8 s) of the 10-s white-noise sound (Figure 1A). Sounds were separated by a 2-s inter-stimulus interval. Participants were informed about whether the gap would occur in the first or the second half of the sound and this expectation was never violated. Participants could thus orient their attention in time, but they did not know exactly when, within the cued half of the sound, the gap would occur. Trials in which the gap occurred in the first half and trials in which the gap occurred in the second half of the sound were presented in separate blocks. The number of blocks and the number of trials per block are described below for each experiment separately.

### Behavioral data analysis

For each sound trial, as long as the gap has not yet occurred, the instantaneous probability of the gap’s occurrence within the cued time window (first half; second half) increases monotonically with the passage of time. This is the “hazard rate” (Niemi and Näätänen, 1981; Nobre et al., 2007; Nobre and van Ede, 2018). Response times are expected to be faster for gaps occurring late compared to early within a cued time window (Niemi and Näätänen, 1981; Nobre et al., 2007). This “hazard-rate effect” indicates successful allocation of attention in time (Nobre et al., 2007). Analyzing behavioral data as a function of time within a cued time window allows us to capture whether individuals orient attention in time during listening.

A gap was considered detected (i.e., a hit) if a response was made between 0.1 and 1.2 s after gap onset, and undetected (i.e., a miss) if no button was pressed within that time window. In order to investigate the effect of the passage of time on behavioral performance, behavioral data were analyzed as a function of the gap time within a cued sound half (ranging from 0 to 2.5 s). In detail, hit rates were analyzed by calculating a logistic regression (generalized linear model with a binomial/logit link function; Tune et al., 2021) that relates single-trial responses to gap times. For the response times (hazard-rate effect; Niemi and Näätänen, 1981; Nobre et al., 2007), a linear function was fit to relate single-trial response times to gap times. The logistic regression (hit rate) and linear function (response time) were fit separately for each participant and condition (e.g., participants cued to the first vs second half of the sound). The slope of the fits describes the relation between hit rate / response time and gap time, and was used as the dependent measure.

### EEG/MEG recordings and preprocessing

Details about EEG and MEG recordings and preprocessing are provided for each experiment below. In short, EEG (16 electrodes) and MEG (204 channels) data were high-pass filtered at 0.7 Hz, low-pass filtered at ∼30 Hz, and divided into epochs ranging from –2 to 12 s time-locked to sound onset. Independent components analysis (ICA; Makeig et al., 1996) was used to identify and suppress activity related to blinks, horizontal eye movements, and the heart.

### Wavelet analysis and alpha power

Wavelet coefficients were calculated by convolving single-trial data epochs (−2 to 12 s) with Morlet wavelets. For each participant, trial, and channel, wavelet coefficients were calculated for frequencies ranging from 1 to 20 Hz (in steps of 0.5 for Experiments 1 and 2; in steps of 1 Hz for Experiment 3) and time points ranging from –1.5 to 10 s (in steps of 0.0156 s for Experiments 1 and 2; in steps of 0.024 s for Experiment 3). Single-trial time-frequency power representations were obtained by squaring the magnitude of the complex wavelet coefficients. Time-frequency power was averaged across trials for each cue condition (first half; second half) and baseline corrected using relative change (Capilla et al., 2014; Wilsch et al., 2015; Dimitrijevic et al., 2017; Henry et al., 2017; Fiedler et al., 2021). That is, the mean power preceding sound onset (–1.5 to –0.2 s) was subtracted from the power values at each time point. The resulting values were divided by the mean power in the pre-sound onset time window (–1.5 to –0.2 s).

Auditory alpha activity often covers a frequency band below 10 Hz (Niedermeyer, 1990; Tiihonen et al., 1991; Lehtelä et al., 1997; Obleser et al., 2012; Billig et al., 2019) compared to the traditional occipital alpha activity peaking at about 10 Hz (Berger, 1929). Indeed, in the experiments of the current study, the frequency of alpha activity was significantly lower than 10 Hz (Figure S1). Alpha power was thus averaged across frequencies between 8 and 10 Hz. This lower frequency band also avoids contributions of power suppressions to the average that were occasionally present at higher frequencies (e.g., Figures 4A, 6A/B; small variations in the selected frequency band do not affect the current results). Please note that analyses for the canonical 8–12 Hz frequency band yielded test statistics that did not differ meaningfully from those for the 8–10 Hz frequency band (test statistics for all alpha power analyses for the 8–12 Hz frequency band are provided in the Supplementary Materials).

In order to investigate whether alpha power is sensitive to participants attending to the first versus second half of the sound, we averaged alpha power in the 0.8-s time interval prior to the 2.5 s during which a gap could occur in the two cue conditions (first interval: 1–1.8 s; second interval: 5.5– 6.3 s). Using alpha power as a dependent measure, an rmANOVA was calculated including the within-participants factors Time Interval (first interval; second interval) and Cue (first half; second half). For Experiment 2, the within-participants factor Task Difficulty (easy; difficult) was additionally included. For Experiment 3, we included the additional between-participants factor Age Group (younger; older).

### Statistical analysis

The experiments reported here were not preregistered. Experimental manipulations were within-participants factors and age group in Experiments 3 and S1 was a between-participants factor. Differences between experimental conditions were thus assessed using one-sample t-tests, paired-samples t-tests, and repeated-measures ANOVAs (rmANOVAs). Age-group differences in Experiment 3 (and Experiment S1) were assessed using a between-participants factor in rmANOVAs or independent-samples t-tests. Reporting of statistical results includes test statistic, degrees of freedom, significance level, and effect size. Details about statistical analyses are provided for each experiment separately below. Throughout the manuscript, effect sizes are provided as partial eta squared (η^2^_p_) for ANOVAs and Cohen’s d for t-tests.

## Experiment 1: Investigation of attention in time during listening

### Methods

#### Participants

Twenty-two participants were recruited for Experiment 1 (age range: 18–31 years; median: 20 years; 14 females and 8 males). For two participants, data from two blocks were excluded because recording of behavioral responses malfunctioned.

#### Procedures

Participants listened to 10-s white-noise sounds in six blocks, each containing 40 trials (N=50 for eight participants). Within each block, a gap could occur at one of 40 (50) uniformly spaced times during a 2.5 s window, randomly drawn without replacement. In three of the six blocks, the gap always occurred within a 2.5-s window in the first half (1.8–4.3 s), whereas in the other three blocks, the gap always occurred within the second half (6.3–8.8 s) of the 10-s white-noise sound (Figure 1A). Hence, participants were presented with 120 (150) trials per cue condition. For each block, participants were cued as to whether the gap would occur in the first or the second half of the sound. Blocks with gaps occurring in the first versus second half of the sound alternated within an experimental session and the starting condition alternated across participants.

#### Behavioral analysis

A dependent samples t-test was used to compare overall hit rate and response times between the two cue conditions (first half; second half). The relationship between gap time and behavioral performance was assessed by comparing hit rate/response time slopes against zero using a one-sample t-test. A dependent-samples t-test was calculated to compare slopes between cue conditions.

#### EEG recordings and preprocessing

EEG was recorded at a 1024-Hz sampling rate from 16 electrodes (Ag/Ag–Cl-electrodes; 10-20 placement) and additionally from the left and right mastoids (BioSemi, Amsterdam, The Netherlands; 208 Hz low-pass filter). Electrodes were referenced online to a monopolar reference feedback loop connecting a driven passive sensor and a common mode sense active sensor, both located posterior on the scalp.

Offline data analysis was carried out using MATLAB software (v7.14; MathWorks, Inc.). Line noise (60 Hz) was suppressed using an elliptic filter. Data were re-referenced to the average mastoids, high-pass filtered at a cutoff of 0.7 Hz (2449 points, Hann window), and low-pass filtered at a cutoff of 30 Hz (111 points, Hann window). Data were divided into epochs ranging from –2 to 12 s (time-locked to sound onset) and ICA (runica method, Makeig et al., 1996; logistic infomax algorithm, Bell and Sejnowski, 1995; Fieldtrip implementation Oostenveld et al., 2011) was used to identify and suppress activity related to blinks and horizontal eye movements. Epochs that exceeded a signal change of more than 250 µV for any electrode following ICA were excluded from analyses.

Time-frequency power was calculated for each cue condition (first half; second half) and alpha-power time courses were calculated as the mean power across the 8–10 Hz frequencies (see General Methods). Alpha activity was averaged across a parietal electrode cluster (Pz, P4, P3) because previous work shows that attention-related alpha activity is strongest at parietal sensors/electrodes and that it originates from parietal cortex (van Dijk et al., 2008; Wöstmann et al., 2015; Henry et al., 2017;

Heideman et al., 2018; Leenders et al., 2018; see also Experiment 3). We averaged alpha power in the 0.8-s time interval prior to the 2.5 s during which a gap could occur in the two cue conditions (first interval: 1–1.8 s; second interval: 5.5–6.3 s). An rmANOVA was calculated using alpha power as the dependent measure and the within-participants factors Time Interval (first interval; second interval) and Cue (first half; second half).

### Results and discussion

Gap-detection hit rates were generally higher (t_21_ = 3.045, p = 0.006, d = 0.649) and response times generally faster (t_21_ = 2.237, p = 0.036, d = 0.477) for gaps occurring in the first compared to the second half of the sound (Figure 1B). This suggests that focusing attention to later parts of the sound is more difficult than focusing on earlier parts. Listeners may struggle more to precisely focus attention when gaps occur in the second half of the sound due to well-known higher variability in time-interval estimations for longer relative to shorter intervals (Weber’s law; Gibbon, 1977; Buhusi and Meck, 2005; Grondin, 2014).

Critically, gap-detection hit rates were higher (t_21_ = 3.746, p = 0.001, d = 0.799) and response times faster (t_21_ = 7.115, p = 5.11 × 10^−7^, d = 1.517) for gaps that occurred later compared to earlier within the cued 2.5-s window (averaged across attention conditions). This was true regardless of whether that window was in the first or second half of the trial (see slopes in Figure 1B). The faster response times for gaps occurring later within a cued half indicate that participants successfully oriented attention in time (hazard-rate effect; Niemi and Näätänen, 1981; Nobre et al., 2007). This response-time effect was smaller when participants attended to the second compared to the first half of the sound (t_21_ = 4.093, p = 5.2 × 10^−4^, d = 0.873). The reduced hazard-rate effect suggests that individuals’ estimation of when a gap may occur is reduced when gaps occur in the second compared to the first half of the sound. This, again, is consistent with higher variability in time estimations for longer relative to shorter intervals (Gibbon, 1977; Buhusi and Meck, 2005; Grondin, 2014).

Alpha power was sensitive to whether participants attended to the first versus second half of the sound: alpha power increased for longer when participants knew the gap would occur later, that is, in the second compared to the first half (Figure 2A/B). For the rmANOVA, we observed a Time Interval × Cue interaction (F_1,21_ = 14.389, p = 0.001, η_p_^2^ = 0.407): Alpha power was larger in the first interval (1–1.8 s) compared to the second interval (5.5–6.3 s) when participants were cued to the first half (F_1,21_ = 7.549, p = 0.012, η_p_^2^ = 0.264), but alpha power was larger in the second interval (5.5–6.3 s) compared to the first interval (1–1.8 s) when participants were cued to the second half of the sound (F_1,21_ = 13.176, p = 0.002, η_p_ ^2^ = 0.386; Figure 2B, bottom). In fact, alpha power increased throughout a trial up to gap occurrence and decreased thereafter (Figure 2C).

**Figure 2:**
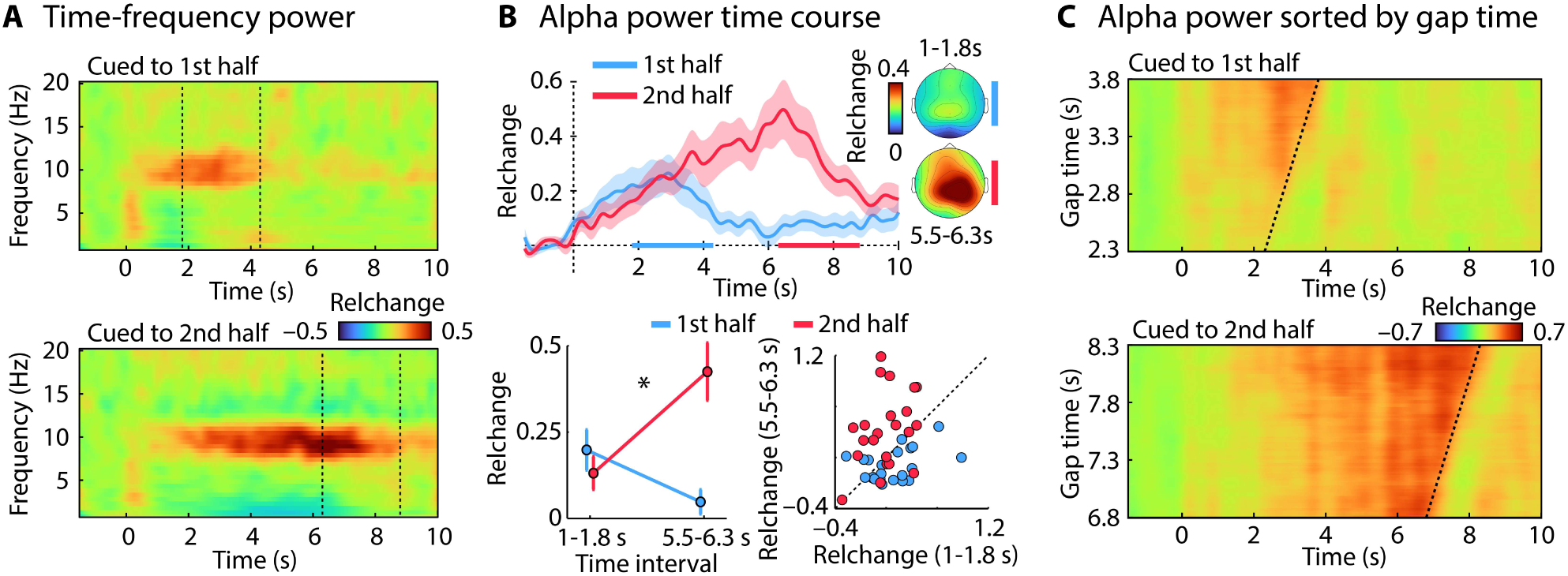
Sensitivity of alpha power to attentive listening in time. A: Time-frequency power (averaged across parietal electrodes; dashed lines mark the 2.5-s windows [gap occurrence]). B: Alpha-power (8–10 Hz) time course for the two cue conditions (first half; second half). Colored lines on the x-axis mark the 2.5-s windows (gap occurrence). Mean alpha power in the 0.8-s time interval prior to the 2.5-s gap window (bottom). C: Alpha-power time courses sorted by gap occurrence. The dashed line indicates the gap onset. Error bars and shadings reflect the standard error of the mean. *p ≤ 0.05, n.s. – not significant.

The results from Experiment 1 show that individuals attend in time during listening according to when they expect a target, and alpha power indicates when in time individuals listen attentively. Our core paradigm is thus suitable for the study of attention regulation in time during listening.

## Experiment 2: Attention regulation at different listening-difficulty levels

Next, we turned to the observation that older adults often experience listening challenges in their everyday lives (Gatehouse and Noble, 2004; Pichora-Fuller et al., 2016; Strauss and Francis, 2017; Peelle, 2018). Individuals can conserve mental resources associated with listening challenges by regulating how much attention is deployed based on the expected difficulty of a listening situation (Brehm and Self, 1989; Richter et al., 2016; Herrmann and Johnsrude, 2020). Alpha power should be sensitive to such regulations of attention based on expected listening difficulty if it is to be useful to the community, but this has not been investigated directly. Experiment 2 aims to investigate how cuing a listener to attend to specific points in time and task difficulty affect behavior and neural alpha activity.

### Methods

#### Participants

Experiment 2 included 20 younger adults (age range: 17–30 years; median: 18; 16 female and 4 males).

#### Procedures

Similar to Experiment 1, we manipulated when a gap occurred (first half; second half), and the cue given prior to a block of trials, in order to manipulate attention in time. In addition, task difficulty was manipulated by changing the duration of the gap (Figure 3A). The estimated gap-duration threshold from the training blocks was used for difficult, short-duration trials (as in Experiment 1), whereas three times the estimated threshold was used for easy, long-duration trials. The different cue (first half; second half) and task-difficulty (easy; difficult) trials were presented in separate blocks. Participants were cued prior to each block whether the gap would occur in the first or the second half of the sound, and whether gap detection would be easy or difficult, and these expectations were never violated. Participants thus had prior knowledge about which sound conditions they would encounter so that they could regulate their attention. Participants listened to 35 trials per block and each of the four block types was presented twice. Within each block, a gap could occur at one of 35 uniformly spaced times during the cued 2.5 s window (cued to first half: 1.8-4.3 s; cued to second half: 6.3-8.8 s), randomly drawn without replacement. Block order was organized such that cue conditions alternated (first half; second half), whereas task-difficulty conditions (easy; difficult) were presented in two consecutive blocks and alternated to create a balanced design. Starting conditions alternated across participants.

**Figure 3:**
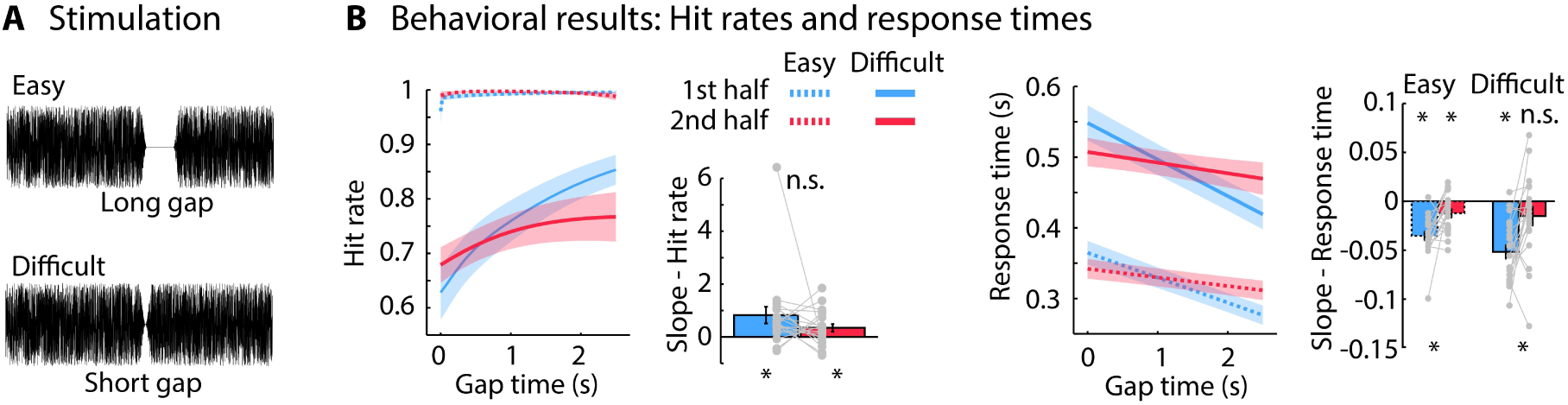
Stimulus representation and behavioral performance for Experiment 2. A: Schematic representation of the gap duration for easy and difficult trials (long and short gap durations, respectively). The duration of the short gap was titrated for each individual to approximately 80%detection rate. The duration of the long gap was three times the duration of the short gap. B: Predicted hit rates and response times from logistic and linear fits, respectively, as a function of gap occurrence within the 2.5-s window, for both cue (first half; second half) and task-difficulty (easy [dashed]; difficult [solid]) conditions. The x-axis reflects the gap time relative to the onset of the cued time window (1.8 s and 6.3 s for first half and second half, respectively). Slopes from logistic (hit rate) and linear (response time) fits are plotted on the right. Slope for hit rates from the easy condition were less reliable due to ceiling effects; they are thus not displayed. Error bars and shadings reflect the standard error of the mean. *p ≤ 0.05, n.s. – not significant.

#### Behavioral data analysis

A rmANOVA was used to compare overall hit rate and response times between the two cue conditions (first half; second half) and the two task-difficulty conditions (easy; difficult). The relationship between gap time and behavioral performance was assessed by comparing hit rate/response time slopes against zero using a one-sample t-test. A rmANOVA and t-tests were used to compare slopes between the cue conditions (first half; second half) and the task-difficulty conditions (easy; difficult).

#### EEG recording and analysis

EEG recording and preprocessing was conducted as for Experiment 1.

Alpha power (8–10 Hz) was averaged in the 0.8-s time interval prior to the cued 2.5-s time window during which a gap could occur in the two cue conditions (first interval: 1–1.8 s; second interval: 5.5–6.3 s), separately for the two cue and task-difficulty conditions. An rmANOVA was calculated using alpha power as the dependent measure and the within-participants factors Time Interval (first interval; second interval), Cue (first half; second half), and Task Difficulty (easy; difficult).

### Results and discussion

Figure 3B depicts the behavioral results for Experiment 2. Hit rates were overall higher (F_1,19_ = 63.007, p = 1.89 × 10^−7^, η_p_^2^ = 0.768) and response times faster (F_1,19_ = 262.594, p = 1.4 × 10^−12^, η_p_^2^ = 0.933) for easy compared to difficult trials, but did not differ between cue conditions (p > 0.25).

Because hit rates for easy trials were at ceiling, the relationship between hit rate and gap time was not analyzed for easy trials. For difficult trials (short gap), hit rates increased with increasing gap time within the cued 2.5-s windows (t_19_ = 3.598, p = 0.002, d = 0.804; averaged across conditions; slopes in Figure 3B). There was no difference between attention conditions (p > 0.2).

Response times decreased with increasing gap time within the cued 2.5-s window (i.e., hazard-rate effect: t_19_ = 7.698, p = 2.96 × 10^−7^, d = 1.721; averaged across conditions; Figure 3B), but this effect was larger when participants attended to the first half compared to the second half of the noise (F_1,19_ = 25.874, p = 6.55 × 10^−5^, η_p_^2^ = 0.577). The hazard-rate effect was unaffected by task difficulty (p > 0.15), suggesting that participants orient attention in time to the same extent for easy and difficult trials (Figure 3B).

Time-frequency power representations and alpha power time courses are depicted in Figure 4A and B, respectively. The rmANOVA for alpha power revealed a Time Interval × Cue interaction (F_1,19_ = 21.001, p = 2.03 × 10^−4^, η_p_^2^ = 0.265). Alpha power was larger in the first interval (1–1.8 s) compared to the second interval (5.5–6.3 s) when participants were cued to the first half (F_1,19_ = 7.697, p = 0.012, η_p_^2^ = 0.288), but alpha power was larger in the second interval (5.5–6.3 s) compared to the first interval (1– 1.8 s) when participants were cued to the second half (F_1,19_ = 21.556, p = 1.8^e-4^, η_p_^2^ = 0.532; Figure 4C). These results replicate the results from Experiment 1 by showing that alpha power indicates when in time individuals listen attentively.

**Figure 4:**
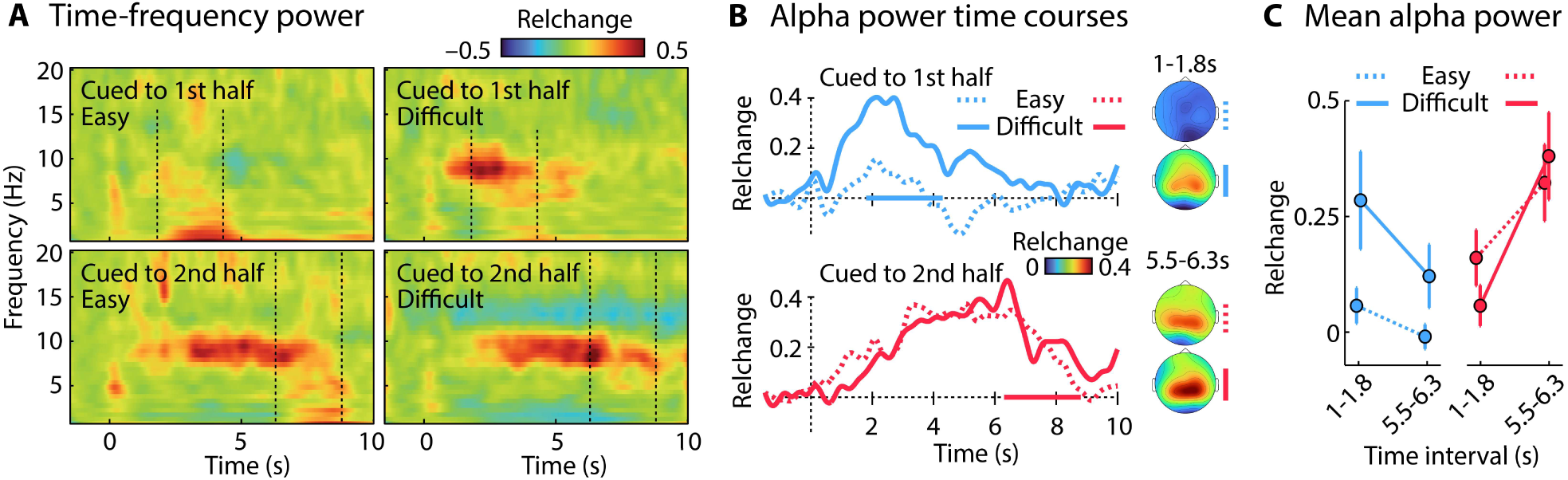
Sensitivity of alpha power to attentional orienting and task difficulty during listening. A: Time-frequency (averaged across parietal electrodes) power representations for each of the four Cue × Task Difficulty conditions. B: Alpha power (8–10 Hz) time courses and topographical distributions. C: Mean alpha power in the 0.8-s time interval prior to the 2.5-s gap windows. Error bars and shadings reflect the standard error of the mean. *p ≤ 0.05, n.s. – not significant.

In addition, the Time Interval × Cue × Task Difficulty interaction was marginally significant (F_1,19_ = 4.022, p = 0.059, η_p_^2^ = 0.175): we conducted an exploratory analysis to understand this marginal interaction in more detail. Separately for each cue condition (first half; second half), a rmANOVA with Time Interval (first interval: 1–1.8 s; second interval: 5.5–6.3 s) and Task Difficulty (easy; difficult) factors was conducted. When participants were cued to the first half of the sound, alpha power was larger for difficult compared to easy trials (effect of Task Difficulty: F_1,19_ = 4.980, p = 0.038, η_p_^2^ = 0.208), but there was no Time Interval × Task Difficulty interaction (p = 0.204). In contrast, when participants were cued to the second half of the sound, alpha power increased over time in both difficulty conditions, but the increase was steeper for difficult compared to easy trials (Time Interval × Task Difficulty interaction: F_1,19_ = 4.638, p = 0.044, η_p_^2^ = 0.196).

Experiment 2 shows that alpha power increased with increasing task difficulty when individuals attended to the first half of the sound. When participants attended to the second half of the sound, alpha power increased over time for both easy- and difficult-to-detect gaps, but the rate of alpha power increase over time was larger. That task difficult had a clearer impact on alpha power when gaps occurred in the first compared to the second half of the sound may be related to overall difficulty differences: Gap detection appears to be easier when gaps occur in the first half compared to the second half of a sound (e.g., Figure 1B, 5A), possibly due to higher variability in time estimations for longer compared to shorter intervals (Gibbon, 1977; Wearden, 2003; Buhusi and Meck, 2005; Grondin, 2014). Alpha power may thus increase when participants attend to the second half of the sound because orienting and maintaining attention over a longer period may be difficult, independent of the experimentally manipulated difficulty (gap duration).

The results from Experiment 2, together with those from Experiment 1, indicate that alpha power is sensitive both to ‘when’ an individual is listening, as well as to ‘how hard’ they are listening (how effortful it is). Alpha power may therefore be a useful way to measure whether younger and older adults differ in how they direct attention to points in time when key information is expected to occur during challenging listening.

## Experiment 3: Assessing age differences in attention regulation in time

Experiment 3 aims to investigate whether behavioral and neural indices of attention regulation in time differ between younger and older adults, and to characterize the brain regions underlying such attention regulation. MEG typically has higher sensitivity, relative to EEG, for the identification of the sources underlying neural responses observed outside of the head, because magnetic fields originating from the brain are less distorted by the skull and scalp than electric potentials (Hämäläinen et al., 1993; Hämäläinen and Hari, 2002; Hari and Puce, 2017). Hence, in order to identify the sources underlying alpha power changes during attention in time, we recorded MEG in younger and older adults using the same experimental procedures as those in Experiment 1 (Figure 1A).

### Methods

#### Participants

Twenty-five younger (age range: 20–33 years; median: 26 years; 15 females and 10 males) and twenty-seven middle-aged to older adults (age range: 54–72 years; median: 63 years; 13 female and 14 males; henceforth referred to as ‘older’ adults) participated in Experiment 3.

Self-rated hearing abilities assessed using the Speech, Spatial and Qualities of Hearing Scale (SSQ; Gatehouse and Noble, 2004) did not differ between age groups (t_49_ = 0.605; p = 0.548; no SSQ ratings were available for one younger participant). Hearing thresholds for the white-noise sound used in the current study were about 6.5 dB higher for older adults compared to younger adults (t_50_ = 5.539, p = 1.1 × 10^−6^). A threshold difference of 6.5 dB is in line with previous studies that compared younger and older adults with clinically ‘normal’ audiometry thresholds (Herrmann et al., 2018, 2022b, a). Finally, to account for any potential differences due to subclinical hearing impairment, sounds were presented at 45 dB above the individual hearing threshold (equating for audibility) and the gap duration was titrated for each participant (equating for task performance/difficulty).

None of the participants showed an indication of cognitive impairment as assessed using the 6-Item Cognitive Impairment Test (6CIT; Upadhyaya et al., 2010; Hessler et al., 2017; younger adults: median score: 0, range: 0-2; older adults: median score: 0, range: 0-4).

#### Procedures

The same experimental procedures as those in Experiment 1 were used (Figure 1A). Gap duration was titrated for each participant so that approximately 80% of gaps were detected successfully. Older adults required the gap to be longer compared to younger adults (t_50_ = 4.282, p = 8.38 × 10^−5^, d = 1.189; mean gap duration and range: younger: 6.72 ms [5.5–8.1 ms], older: 7.87 ms [6–9.9 ms]) in order to achieve a similar detection rate, consistent with previous gap-detection observations (Snell, 1997; Strouse et al., 1998; Humes et al., 2009; Fitzgibbons and Gordon-Salant, 2010).

#### Behavioral data analysis

A rmANOVA was used to compare overall hit rate and response times between the two cue conditions (within-participants factor: first half; second half) and the two age groups (between-participants factor: younger; older). The relationship between gap time and behavioral performance was assessed by comparing hit rate/response time slopes against zero using a one-sample t-test. A rmANOVA and t-tests were used to compare slopes between cue conditions (first half; second half) and age groups (younger; older).

#### MEG recordings and analysis

Magnetoencephalographic data were recorded using a 306-channel Neuromag Vectorview MEG (Elekta; sampling rate: 1000 Hz, filter: DC–330 Hz) at the Max Planck Institute for Human Cognitive and Brain Sciences in Leipzig, Germany. The signal space separation (SSS) method was used to suppress external interference, interpolate bad channels, and transform individual data to a common sensor space. The common sensor space enables comparison of topographical distributions across participants (Taulu et al., 2004; Taulu et al., 2005). We focused on the 204 orthogonal planar gradiometers in 102 locations, because they are most sensitive to magnetic fields originating from sources directly below them (Hämäläinen et al., 1993). Note that the information in magnetometers and gradiometers after SSS correction carry redundant information and thus no information is lost by focusing on gradiometers (Garcés et al., 2017; Herrmann et al., 2018).

Data were high-pass filtered (0.7 Hz; 2391 points, Hann window), low-pass filtered (33.3 Hz, 101 points, Hann window), down-sampled to 500 Hz, and divided into epochs ranging from –2 s to 12 s time locked to sound onset. Independent components analysis (Makeig et al., 1996; Oostenveld et al., 2011) was used to remove noisy channels and activity related to blinks, eye movements, and the heart. Epochs comprising a signal change larger than 300 pT/m in any gradiometer channel following ICA were excluded.

For each participant, time-frequency power was calculated for each cue condition (first half; second half). Alpha-power time courses were calculated by averaging power across the 8 to 10 Hz frequency window. Alpha power (8–10 Hz) was averaged in the 0.8-s time interval prior to the 2.5 s during which a gap could occur in the two cue conditions (first interval: 1–1.8 s; second interval: 5.5–6.3 s). A rmANOVA with within-participants factors Time Interval (first interval; second interval) and Cue (first half; second half), and the between-participants factor Age Group (younger; older) was calculated.

#### MEG source reconstruction

Anatomically constrained source localization was used in order to identify the neural sources underlying alpha power. Individual T1-weighted MR images (3T Magnetom Trio, Siemens AG, Germany) were used to construct inner-skull surfaces (volume conductor) and mid-gray matter cortical surfaces (source model; Freesurfer and MNE software). Inner-skull surfaces consisted of a mesh of consisted of a mesh of 2,562 vertices and cortical surfaces of 10,242 vertices per hemisphere. No MRI scan was available for one older participant and data from this person were thus not considered for source analyses and related statistical analyses. The MR and the MEG coordinate systems were co-registered using MNE software which included an automated and iterative procedure that fitted the >300 digitized head surface points (Polhemus FASTRAK 3D digitizer) to the MR reconstructed head surface (Besl and McKay, 1992). For each participant, a boundary element model (inner skull) was constructed using MNE software and a lead field was calculated using Fieldtrip (Nolte, 2003).

Source reconstructions for time-frequency representations were calculated using dynamic imaging of coherent sources (DICS; Gross et al., 2001) with custom MATLAB scripts. DICS is a beamforming technique that involves estimating a spatial filter at each source location. The cross-spectral density matrix was calculated from the complex wavelet coefficients of all trials in the 8–10 Hz frequency window and 0.8–9.2 s time window, and used to estimate real-numbered spatial filters (using the dominant direction; Gross et al., 2001). The 0.8–9.2 s time window was chosen to ensure that power related to onset or offset responses do not influence the filter calculation. Single-trial wavelet coefficients were projected into source space using these filters and power time courses were calculated similar to the sensor space analysis.

Individual cortical representations were transformed to a common coordinate system (Fischl et al., 1999b) in order to enable region of interest identification based on standard anatomical parcellations (Glasser et al., 2016). To this end, neural activity was mapped to each participant’s full Freesurfer in source space mesh (>100,000 vertices per hemisphere), spatially smoothed across the surface using an approximation to a 6-mm FWHM Gaussian kernel (Han et al., 2006), and morphed to the standard Freesurfer cortical surface (Fischl et al., 1999a).

#### Statistical analysis of source data

Analysis of age differences in spatial location of alpha activity focused on a pre-gap time window independent of cue conditions. To this end, alpha power was averaged in the −0.8 to −0.2 s time window prior to each trial’s specific gap onset (−0.2 was chosen to avoid that gap-related activity influences pre-gap alpha power). Age-differences in spatial configuration of alpha activity were assessed by calculating for each participant and brain hemisphere the 3D coordinate at which pre-gap alpha power was maximal. Independent-samples t-tests were used to compare the x, y, and z coordinate between age groups.

Analysis of inter-subject correlation of alpha power maps was additionally used to examine age differences in spatial configuration of alpha activity. Each participant’s spatial map of pre-gap alpha power was separately correlated with the spatial map of every other participant (vertices were used as samples). For each participant, this resulted in 50 correlation values. The 50 correlation values were categorized into two categories: correlations with members of the same age group (within) and correlations with members of the other age group (between). For each participant, the average correlation value was calculated for each of the two categories. This resulted in two mean correlation values for each participant, reflecting the degree of similarity of the activation map to individuals within the same versus the other age group. If spatial maps differ between age groups, we would expect larger correlations with within-group than between-group members. We used independent-samples t-tests to test this.

Previous work suggests task-related alpha activity originates from sensory cortices and parietal cortex (Tuladhar et al., 2007; van Dijk et al., 2008; Müller et al., 2015; Wöstmann et al., 2016; Tune et al., 2018). In order to analyze age differences in attentional modulation of alpha power, we used Glasser’s anatomical parcellation of the cortical surface (Glasser et al., 2016) to obtain vertices for bilateral superior temporal cortex (STC) and bilateral superior parietal cortex (SPC; Figure 6). Alpha power was averaged within each region, resulting in region-specific alpha-power time courses. Alpha power was averaged in the 0.8-s interval prior to the 2.5 s during which a gap could occur (first interval: 1–1.8 s; second interval: 5.5–6.3 s). An rmANOVA was calculated using the within-participants factors Time Interval (first interval; second interval), Cue (first half; second half), and Region (STC, SPC), and the between-participants factor Age Group (younger; older).

### Results and discussion

#### Age differences in behavioural accuracy but not attentional orienting

Analysis of hit rates revealed no overall difference between age groups (F_1,50_ = 1.457, p = 0.233, η_p_^2^ = 0.028). Gap-detection hit rates were generally higher for gaps occurring in the first compared to the second half of the sound (F_1,50_ = 8.312, p = 0.006, η_p_^2^ = 0.143), as in Experiment 1, perhaps because focusing attention on later parts of the sound is more difficult than focusing on earlier parts.

Critically, hit rates increased with increasing passage of time within the cued 2.5 s window for younger people (t_24_ = 3.467, p = 0.002, d = 0.693), but decreased (numerically) for older people (t_26_ = - 0.187, p = 0.853, d = 0.036) as indicated by the age-group difference between slopes (F_1,50_ = 4.817, p = 0.033, η_p_^2^ = 0.088; Figure 5A).

**Figure 5:**
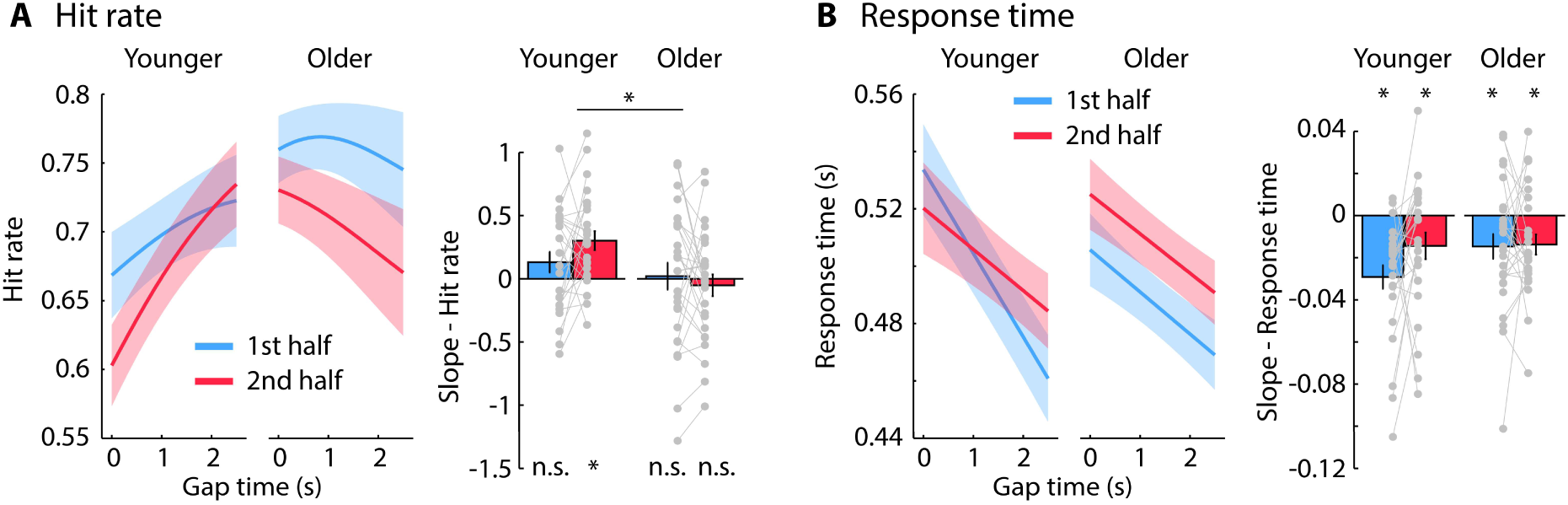
Behavioral results for younger and older adults. Predicted hit rates and response times from logistic and linear fits, respectively, as a function of gap occurrence within the cued 2.5-s window. The x-axis in panels A and B reflects the gap time relative to the onset of the cued time window (1.8 s and 6.3 s for first half and second half, respectively). Error bars and shadings reflect the standard error of the mean. *p ≤ 0.05, n.s. – not significant.

Analysis of response times revealed no overall difference between age groups (F_1,50_ = 0.005, p = 0.944, η_p_^2^ < 0.001). Response times decreased with increasing gap time within a cued 2.5-s time window (i.e., hazard-rate effect; t_51_ = 6.035, p = 1.81 × 10^−6^, d = 0.837; average across cue conditions; Figure 5B). There was no effect of age group or interactions involving age groups on response times, although there was a trend towards a larger hazard-rate effect when younger adults were cued to the first half of the sound compared to the second half (t_24_ = 1.792, p = 0.086, d = 0.358; as for Experiment 1; Figure 1B), whereas the hazard rate effect did not differ between cue conditions for older adults (t_24_ = 0.131, p = 0.896, d = 0.025).

In sum, there were no overall differences in hit rates or response times between younger and older adults, since gap duration was manipulated to achieve a hit rate of ∼80% for each participant. Response-time data indicate that both younger and older adults successfully oriented attention in time during listening, although there was some indication that younger adults may use timing information differently (trending slope differences) compared to older adults. A different picture emerged for hit rates: Younger individuals detected more gaps when they occurred later compared to earlier within a cued sound half, whereas no difference was found for older individuals, potentially suggesting age differences in cognitive control processes (discussed in detail below).

#### Younger and older adults show attention-related alpha power modulations

MEG data were first analyzed in sensor space to examine whether MEG alpha power is sensitive to when participants attend (Figure 6A-D). As for Experiments 1 and 2, the Time Interval × Cue interaction (F_1,50_ = 14.994, p = 3.14 × 10^−4^, η_p_^2^ = 0.231) showed that alpha power was larger in the first interval (1-1.8 s) compared to the second interval (5.5-6.3 s) when participants were cued to the first half (F_1,51_ = 10.646, p = 0.002, η_p_^2^ = 0.173), whereas alpha power was larger in the second compared to the first interval when participants were cued to the second half (F_1,51_ = 12.604, p = 8.4 × 10^−4^, η_p_^2^ = 0.198; Figure 6C,D). This interaction indicates that alpha power was sensitive to whether participants attended to the first versus second half of the sound.

We also observed a Cue × Age Group interaction (F_1,50_ = 5.240, p = 0.026, η_p_^2^ = 0.095): Averaged across the two analysis time intervals (first and second interval), alpha power was numerically larger for younger (p = 0.16) but smaller for older adults (p = 0.085) when they were cued to the first compared to the second half of the sound (Figure 6C/D, right). None of the other effects were significant.

#### Age-differences in spatial distribution of alpha activity

Analyses described in the previous section were based on activity at MEG sensors, but this potentially reflects the summed activity from distinct sources (Hämäläinen et al., 1993; Hämäläinen and Hari, 2002). To gain spatial specificity, we calculated source localizations of alpha power. For younger adults, alpha power was strongest in superior temporal cortex and superior parietal cortex (Figure 6E). For older adults, alpha activity was strongest in superior and posterior temporal cortex, but appeared weaker in superior parietal cortex (Figure 6F; see also Figure S2).

**Figure 6:**
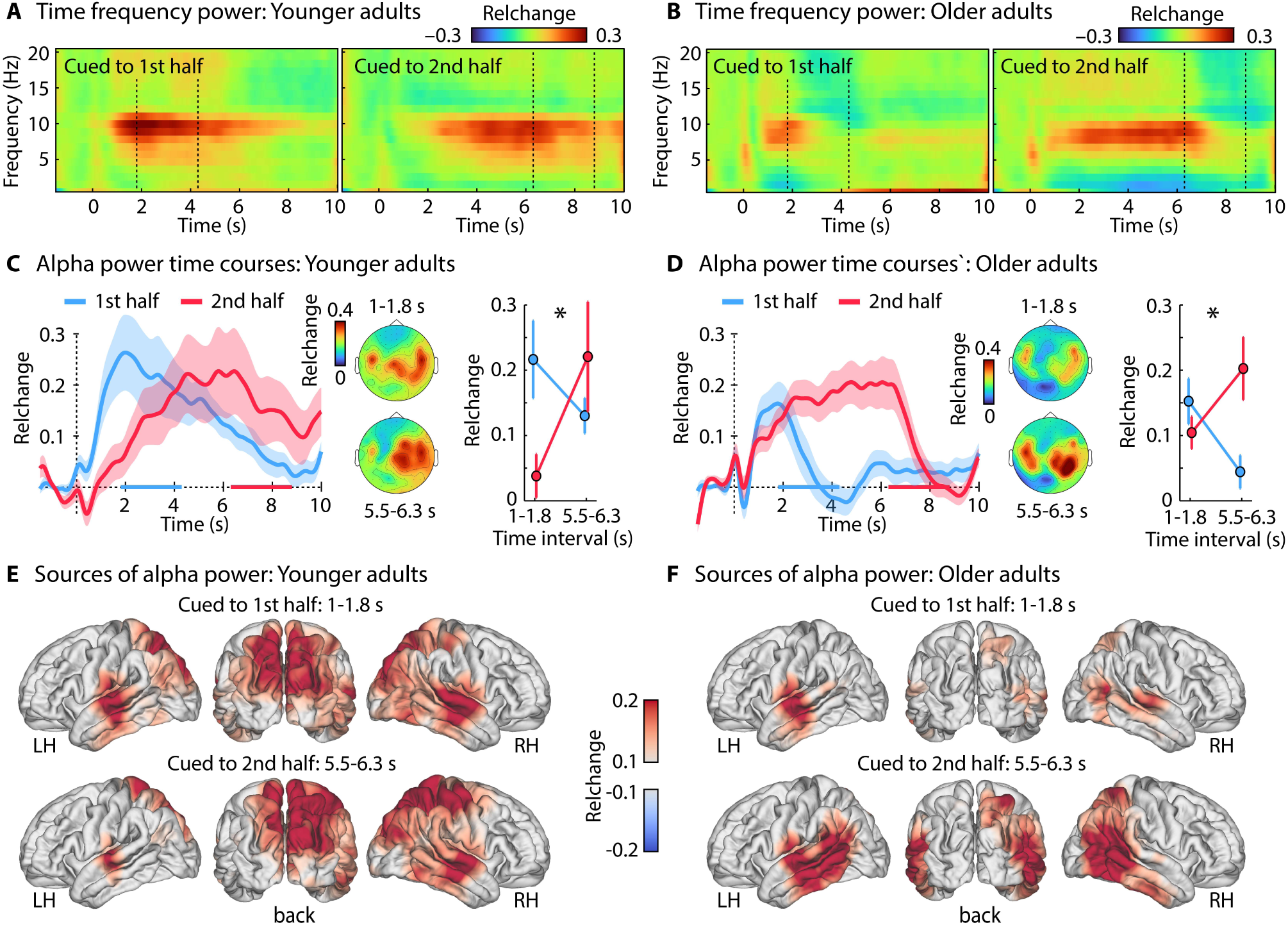
Effect of age group on listening-related alpha power. A: Time-frequency power (averaged across channels; dashed lines mark the 2.5-s gap window). B: Alpha-power (8–10 Hz) time courses for both cue conditions (first half; second half). Colored lines on the x-axis mark the 2.5-s gap window. Topographies and averaged alpha power are shown for the 0.8-s time interval prior to the 2.5 s during which a gap could occur (first interval: 1–1.8 s; second interval: 5.5–6.3 s). The interaction was significant in younger and older adults. C: Source localization of alpha power for both time intervals and age groups. Error bars and shadings reflect the standard error of the mean. *p ≤ 0.05.

The analysis of age-group differences in the spatial configuration of alpha power revealed that peak alpha power was consistently more ventro-lateral in older compared to younger adults (Figure 7A; x-axis of left hemisphere: t_49_ = 2.833, p = 0.007, d = 0.794; x-axis of right hemisphere: t_49_ = 1.880, p = 0.066, d = 0.527; z-axis of left hemisphere: t_49_ = 3.367, p = 0.002, d = 0.943; z-axis of right hemisphere: t_49_ = 2.420, p = 0.019, d = 0.678).

**Figure 7:**
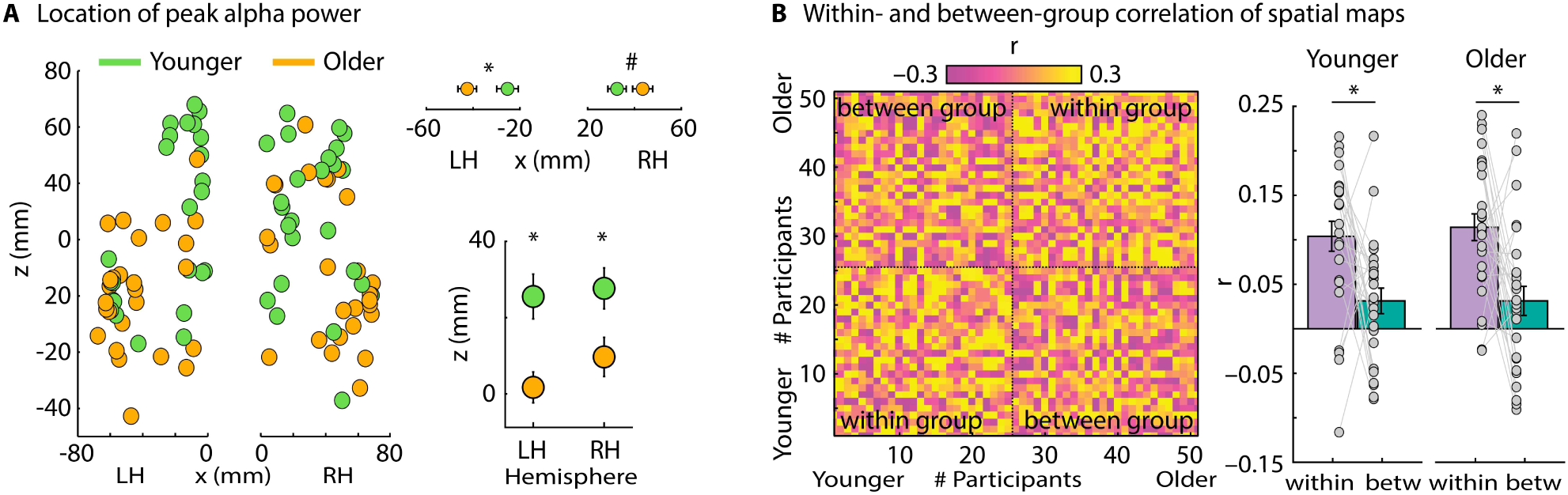
Age-related differences in spatial configuration underlying alpha power. A: Spatial coordinates of alpha power activation peaks for the left (LH) and right hemisphere (RH). Data on the right reflect the mean location on the x-axis (left-right) and z-axis (ventral-dorsal). Activation maxima were more ventro-lateral in older compared to younger adults. B: Pair-wise correlation of spatial maps of alpha power among all participants. Correlations were higher among participants from the same age group (within) than with members from the other age group (between), for both younger and older adults. Error bars reflect the standard error of the mean. *p ≤ 0.05, # ≤ 0.1

Correlations of activation maps among participants further highlight age differences in the spatial configuration of brain activations: Correlations of spatial alpha-power maps were larger among members of the same age group (within) compared to members of the other age group (between; Figure 7B; younger adults: t_24_ = 2.812, p = 0.01, d = 0.562; older adults: t_25_ = 3.243, p = 0.003, d = 0.636). These results suggest that the source configuration underlying listening-related alpha activity differs between younger and older adults.

In Experiment S1 (see Supplementary Materials), we further show that alpha activity during active listening (i.e., when there is a task) originates from spatially distinct brain regions compared to the traditional resting-state alpha activity originating from occipital cortex (Figure S3). This suggests that there are multiple alpha oscillators in the brain (cf., Başar et al., 1997; Klimesch, 1999; Bollimunta et al., 2008; van Dijk et al., 2008; Wöstmann et al., 2020). In contrast to alpha activity during active listening, the sources underlying resting-state alpha activity do not differ between younger and older adults (Supplementary Materials), suggesting age-specific changes in the listening-related alpha oscillator.

#### Attentional modulation of alpha power in superior parietal cortex is reduced in older adults

To investigate the sensitivity of brain regions to attention regulation in time, alpha-power time courses were analyzed for the superior temporal and superior parietal cortical regions (Figure 8).

**Figure 8:**
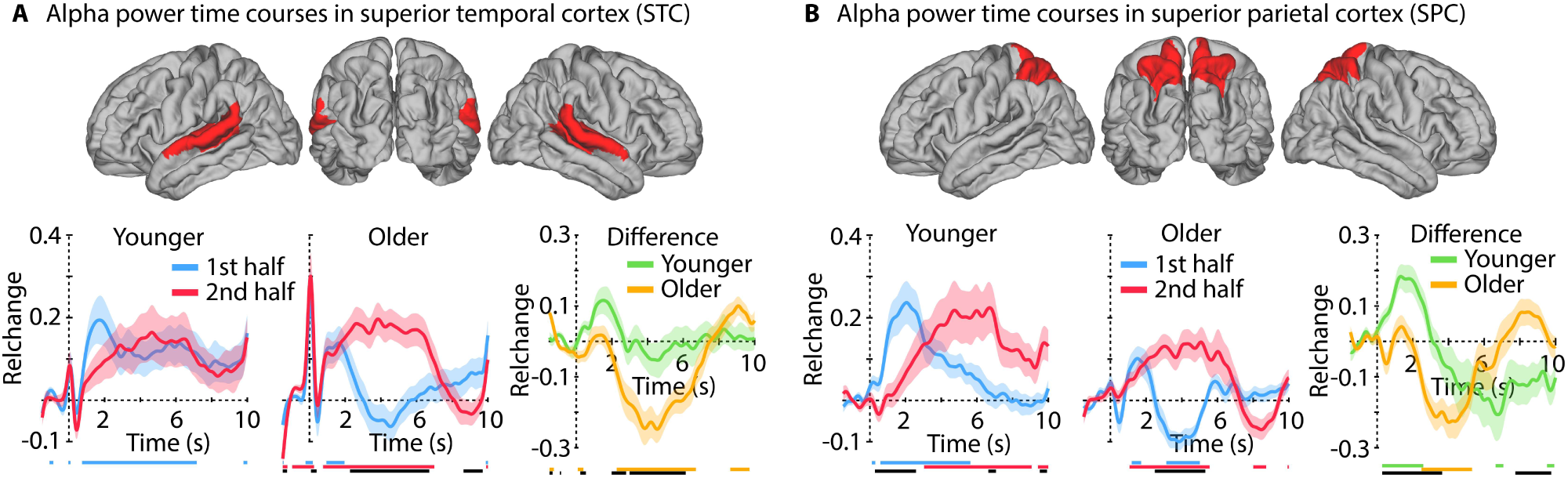
Alpha-power time courses for two anatomically defined regions of interest. A: Superior temporal cortex region. Alpha power time courses for both cue conditions (first half; second half) for both age groups, and time courses for the difference between cue conditions. Colored lines at the bottom indicate a significant difference from zero for the related time course (p ≤ 0.05, FDR-thresholded). The black line indicates a significant difference between time courses. Shadings around mean time courses reflect the standard error of the mean. B: Same as panel A for the superior parietal cortex region.

For younger people, alpha-power differences between cue conditions (first half; second half) were strongest in superior parietal cortex, whereas no difference was observed for superior temporal cortex (Figure 8). For older people, differences between cue conditions were present in both the superior temporal and the superior parietal regions (Figure 8), consistent with the different source configurations in older adults (Figure 6E/F).

To statistically quantify age-group differences in attentional modulation of alpha activity, we conducted an rmANOVA with factors Cue (first half; second half), Time Interval (first interval: 1–1.8 s; second interval: 5.5–6.3 s), Region (STC; SPC), and Age Group (younger; older). The Cue × Time Interval × Region × Age Group interaction was significant (F_1,49_ = 16.935, p = 1.48 × 10^−4^, n_p_^2^ = 0.257; Figure 9A and B). In younger adults, the Cue × Time Interval interaction was significant only for SPC, but not for STC (Cue × Time Interval × Region: F_1,24_ = 18.029, p = 2.83 × 10^−4^, n_p_^2^ = 0.429), showing that alpha power in SPC, but not in STC, decreased from the 1–1.8 s (first interval) to 5.5–6.3 s (second interval) when participants were cued to the first half of the sound, and increased from the 1–1.8 s to 5.5–6.3 s intervals when participants were cued to second half of the sound. In older adults, the Cue × Time Interval interaction (F_1,25_ = 9.885, p = 0.004, n_p_^2^ = 0.283) was not further specified by brain region (Cue × Time Interval × Region: F_1,25_ = 2.466, p = 0.129, n_p_^2^ = 0.09).

**Figure 9:**
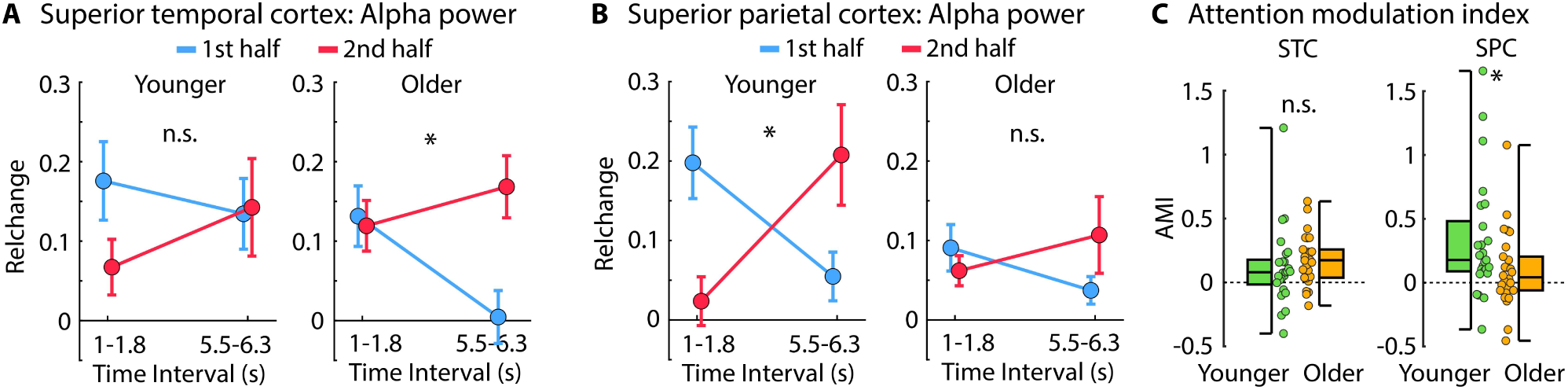
Alpha power in the 0.8-s interval prior to the 2.5 s during which a gap could occur (first interval: 1– 1.8 s; second interval: 5.5–6.3 s). The Cue × Time Interval interaction was greater in the superior parietal cortex for younger compared to older adults, whereas age groups did not differ for the superior temporal cortex (Cue × Time Interval × Region × Age Group interaction: p = 1.48 × 10^−4^; A and B). Panel C shows the interaction as an Attention Modulation Index (AMI). AMI was calculated as the difference between the first and the second interval when participants were cued to first half, plus the difference between the second and the first interval when participants were cued to second half. Larger values mean a stronger modulation of alpha power by the attentional cue. *p ≤ 0.05; n.s. – not significant; STC – superior temporal cortex; SPC – superior parietal cortex

To examine the Cue × Time Interval × Region × Age Group interaction in a different way, we also calculated an attentional modulation index (AMI) as the sum of the difference between time intervals (i.e., difference between first and second interval for cued to first half, plus the difference between second and first interval for cued to second half), separately for each participant and region. The attentional modulation was larger in SPC for younger than older adults (t_49_ = 2.094, p = 0.041, d = 0.587), whereas there was no difference between age groups for STC (t_49_ = −0.829, p = 0.411, d = 0.232; Region × Age-Group interaction: F_1,49_ = 16.935, p = 1.48 × 10^−4^, n_p_^2^ = 0.257; Figure 9C).

In sum, the source localization data indicate age-related differences in the brain regions underlying listening-related alpha activity. Alpha power and alpha-power modulations by attention were largest in superior parietal cortex in younger adults, whereas these effects were reduced in older adults. In contrast, older adults exhibit an activation maximum located more ventro-laterally in the brain, covering posterior temporal cortex. Below, we discuss how this age difference in the brain regions underlying attention regulation may be the result of more proactive (younger) vs reactive (older) cognitive control strategies during challenging listening.

## General discussion

In the current study, we investigated the neural processes involved when individuals listen attentively at specific points in time and the extent to which these processes differ between younger and older adults. We demonstrate that alpha power indicates when in time individuals listen attentively (Experiment 1) and that alpha power is modulated by listening difficulty, taken to reflect how much attention is deployed (Experiment 2). Our data show that older adults are able to regulate attention in time, but that younger and older adults recruit different brain regions during attentive listening. Younger adults show strongest attention-related modulation of alpha power in superior parietal cortex, whereas older adults show strongest modulations in posterior temporal cortex and reduced activity in superior parietal cortex (Experiment 3). We suggest this dissociation in source location between younger and older groups may be the result of different cognitive control strategies employed to regulate attention in time during challenging listening.

### Alpha power is sensitive to critical aspects of attention regulation during listening

We demonstrate that neural alpha power in superior parietal cortex reflects individuals’ directed attention to points in time when key information is expected to occur during challenging listening (Figures 2, 4, 6, and 8). Our work is consistent with studies showing that alpha power aligns temporally with anticipated visual stimuli (Rohenkohl and Nobre, 2011; Bonnefond and Jensen, 2012; Payne et al., 2013). Previous work also suggests that alpha activity is sensitive to temporally dynamic auditory attention, where alpha power synchronizes with attended words presented during dichotic speech listening (Wöstmann et al., 2016; Tune et al., 2018; Wöstmann et al., 2021). We show that alpha power reflects expectations about when key auditory information will occur, thus indexing attention deployment.

Alpha power has been shown to increase when individuals listen attentively to auditory stimuli relative to a baseline, passive listening, or visual stimulation (Adrian, 1944; Foxe et al., 1998; Wöstmann et al., 2015; Wöstmann et al., 2016; Dimitrijevic et al., 2017; Henry et al., 2017; Wöstmann et al., 2019; Fiedler et al., 2021). Alpha power also increases in the hemisphere that is ipsilateral (relative to contralateral) to the side to which a listener attends (Banerjee et al., 2011; Tune et al., 2018; Dahl et al., 2019; Deng et al., 2020; Tune et al., 2021). Our data demonstrate that alpha power also indicates the degree to which a listener deploys attention based on expected listening difficulty (Figure 4). Alpha power was greater when participants expected gap detection to be difficult compared to easy, although this was only observed when participants expected the gap to occur within ∼4 s after sound onset. Estimating the precise timing of temporally more distant events is more difficult (Gibbon, 1977; Wearden, 2003; Buhusi and Meck, 2005; Grondin, 2014). Along these lines, we observed lower overall performance (Figure 1 and 5) and a smaller hazard-rate effect (Figure 1) when participants attended to the second compared to the first half of the sound. High levels of attention may thus have been needed for the detection of gaps in the second half of the sound, independent of whether gap detection was expected to be easy or difficult. This additional load may have led to the increase in alpha power for both difficulty conditions when participants attended to the second half of the sound (Figure 4B/C). Our results are compatible with the idea that attention enables enable listeners to budget mental resources effectively, allocating enough resources moment to moment in order to achieve listening goals.

### Age differences in behavior may reflect different cognitive control strategies

The hazard-rate effect for response times indicates that both age groups oriented attention in time (Figure 5; Nobre et al., 2007; Nobre and van Ede, 2018). Some previous studies have suggested older adults may not use information to orient attention in time as well as younger adults do (Vallesi et al., 2009; Zanto et al., 2011; Heideman et al., 2018), whereas this was not observed in other studies (Chauvin et al., 2016; Droit-Volet et al., 2019). Interpretation of these results is difficult, however, because older adults often responded overall more slowly than younger adults. Here, overall responses times were similar between age groups. The comparable hazard-rate effect in both age groups suggests that older adults are well able to orient attention in time during listening. Gap-detection performance, in contrast, differed between age-groups. Younger individuals detected more gaps when they occurred later, compared to earlier, within each cued sound half, whereas older individuals detected fewer (although for older adults the decline was not significant; Figure 5).

The cognitive-control literature distinguishes between ‘proactive’ and ‘reactive’ control (Braver et al., 2007; Braver, 2012). Proactive cognitive control is anticipatory and refers to the active and sustained maintenance of goals and related information to facilitate task performance. In contrast, reactive cognitive control is not anticipatory: goals and contextual information are only transiently activated and decay faster (Braver et al., 2007). Behavioral performance in tasks requiring attentional orienting will thus depend on the cognitive control strategy employed (Braver et al., 2007). Critically, proactive control may be impaired in older adults, whereas reactive control has been suggested to remain intact in older adulthood (Braver et al., 2007; Braver, 2012). In the current listening task, decreasing hit rates for later gaps within a cued sound half for older adults may indicate a decay in maintenance of goal and context information reflective of more reactive strategy.

Our data provide other indications that temporal information is somewhat less anticipatorily utilized by older adults. Whereas response times depended on when a gap occurred in the first compared to the second half of the sound in younger adults (Figures 1B and 5B), this difference was clearly absent in older adults (Figure 5B). In fact, the influence of gap time on response times was very similar for younger adults cued to the second sound half and those for older adults cued to either sound half (Figure 5B, the decrease in response times with gap time for older adults’ was also smaller compared to the response-time decrease for younger adults cued to the first half; for all p < 0.05, one-sided). This may suggest that older adults, while orienting attention in time, may have done so less effectively. We speculate that the pattern of hit rates and responses times is consistent with older adults employing a mixture of proactive and reactive cognitive control strategies, compared to younger adults, who relied more on a proactive strategy.

### Older adults recruit different brain regions for attention regulation than younger adults

Consistent with previous reports, we show an age-related reduction of alpha activity in superior parietal cortex (Figure S2; Sander et al., 2012; Vaden et al., 2012; Deiber et al., 2013; Hong et al., 2015; Wöstmann et al., 2015; Henry et al., 2017; Leenders et al., 2018; Rogers et al., 2018; Bonacci et al., 2019; Getzmann et al., 2020), suggesting that parietal cortex is less involved in challenging attention tasks in older adults.

Alpha power reflected attentional orienting in time for both younger and older adults (Figures 6, 8), but the recruited brain regions differed between age groups. Alpha activity in superior parietal cortex was modulated by attention in younger adults (Tuladhar et al., 2007; van Dijk et al., 2008; Tune et al., 2018), whereas older adults showed attention-related modulations in posterior temporal cortex (Figures 6, 8, 9). Previous studies using visuo-spatial attention tasks have reported reduced attention-related lateralization of parietal alpha power for older compared to younger adults (Sander et al., 2012; Deiber et al., 2013; Hong et al., 2015; Mok et al., 2016; Heideman et al., 2018; Leenders et al., 2018; Getzmann et al., 2020). Source localization was not conducted in these previous studies, which may therefore have missed brain regions sensitive to attention regulation in older adults. Other research in the auditory domain shows the presence of alpha-power lateralization in a spatial-attention task in middle-aged to older adults, but comparisons with younger individuals were not conducted (Tune et al., 2018; Tune et al., 2021).

Proactive cognitive control is associated with sustained activity in prefrontal cortex (Braver et al., 2007; Braver, 2012). Prefrontal cortex, in turn, is thought to orchestrate other task-relevant regions by providing representation and maintenance of contextual information, such as the attention cue used here (Braver et al., 2007). Parietal cortex is crucial for performance of tasks with attentional load (Kanwisher and Wojciulik, 2000; Pessoa et al., 2003; Braver, 2012; Power and Petersen, 2013; Zanto and Gazzaley, 2017) and may, in younger adults, reflect the outcome of cognitive control processes in prefrontal cortex. Prefrontal activity was not observed directly here, perhaps because alpha oscillators are commonly associated with parietal and sensory cortices (Başar et al., 1997; van Dijk et al., 2008; Wöstmann et al., 2016), making our measures more sensitive to regions over which prefrontal cortex exhibits control than to prefrontal regions themselves. Reactive control can involve other brain regions, such as anterior cingulate or other more posterior cortices, including temporal cortex, to mediate performance in challenging tasks (Speer et al., 2003; Braver et al., 2007; Dew et al., 2011; Braver, 2012). Recruitment of different brain regions in older compared to younger adults in the current study could thus indicate more reliance on reactive control processes.

## Conclusions

We investigated the neural processes involved when individuals listen attentively at specific points in time and asked whether these processes differ between younger and older adults. Alpha power reflected *when in time* individuals listened attentively and *how much they attended* (induced by listening difficulty). Critically, our data show that older adults are able to regulate attention in time during listening, but that they appear to use different behavioral listening strategies and recruit different brain regions than do younger adults. Younger adults show attention-related modulations of alpha power in superior parietal cortex, whereas older adults show attention-related modulation in posterior temporal cortex. We suggest that the observed age-group differences in recruited neural circuits and behavior reflect different cognitive control strategies employed to regulate attention in time during challenging listening.

## Acknowledgements

Research was supported by the Canadian Institutes of Health Research (MOP133450 to I.S. Johnsrude), the Natural Sciences and Engineering Research Council of Canada (RGPIN-2021-02602 to BH), and the German Research Foundation (Deutsche Forschungsgesellschaft; DFG; HE 7857/1-1 to BH). BH was supported by a BrainsCAN Tier I postdoctoral fellowship (Canada First Research Excellence Fund; CFREF) and the Canada Research Chairs Program (232733). We thank the Max Planck Institute for Human Cognitive and Brain Sciences for the opportunity to record the MEG data and Yvonne Wolff-Rosier for help during data acquisition.

## Conflict of Interest

The authors have no conflict of interest.

## Author Contributions

BH conceptualized and designed the research, recorded and analyzed the data, and wrote the manuscript. BM assisted with MEG data acquisition and MEG data analysis, and wrote the manuscript. MJH and JO provided conceptual support and wrote the manuscript. ISJ conceptualized and designed the research and wrote the manuscript.

## Notes

### Competing Interest Statement

The authors have declared no competing interest.

### Summary of Updates

Added hearing and cognitive assessment data. Minor revisions in abstract and main text.

## References

Adrian ED (1944) Brain rhythms. Nature 153:360–362.

Banerjee S, Snyder AC, Molholm S, Foxe JJ (2011) Oscillatory Alpha-Band Mechanisms and the Deployment of Spatial Attention to Anticipated Auditory and Visual Target Locations: Supramodal or Sensory-Specific Control Mechanisms? The Journal of Neuroscience 31:9923–9932.

Başar E, Schürmann M, Başar-Eroglu C, Karakaş S (1997) Alpha oscillations in brain functioning: an integrative theory. International Journal of Psychophysiology 26:5–29.

Bell AJ, Sejnowski TJ (1995) An information maximization approach to blind separation and blind deconvolution. Neural Computation 7:1129–1159.

Berger H (1929) Über das Elektrenkephalogramm des Menschen. European Archives of Psychiatry and Clinical Neuroscience 87:527–570.

Besl PJ, McKay ND (1992) A method for registration of 3-D shapes. IEEE Transactions On Pattern Analysis And Machine Intelligence 14:239–256.

Billig AJ, Herrmann B, Rhone AE, Gander PE, Nourski KV, Snoad BF, Kovach CK, Kawasaki H, Howard III MA, Johnsrude IS (2019) A sound-sensitive source of alpha oscillations in human non-primary auditory cortex. The Journal of Neuroscience 39:8679–8689.

Bollimunta A, Cheng Y, Schroeder CE, Ding M (2008) Neuronal Mechanisms of Cortical Alpha Oscillations in Awake-Behaving Macaques. The Journal of Neuroscience 28:9976–9988.

Bonacci LM, Dai L, Shinn-Cunningham BG (2019) Weak neural signatures of spatial selective auditory attention in hearing-impaired listeners. The Journal of the Acoustical Society of America 146:2577–2589.

Bonnefond M, Jensen O (2012) Alpha Oscillations Serve to Protect Working Memory Maintenance against Anticipated Distracters. Current Biology 22:1969–1974.

Braver TS (2012) The variable nature of cognitive control: a dual mechanisms framework. Trends in Cognitive Sciences 16:106–113.

Braver TS, Barch DM (2002) A theory of cognitive control, aging cognition, and neuromodulation. Neuroscience and Biobehavioral Reviews 26:809–817.

Braver TS, Gray JR, Burgess GC (2007) Explaining the many varieties of working memory variation: Dual mechanisms of cognitive control. In: Variation in working memory (Conway A, Jarrold C, Kane M, Miyake A, Towse J, eds), pp 76–106. New York, NY, US: Oxford University Press.

Brehm JW, Self EA (1989) The Intensity of Motivation. Annual Review of Psychology 40:109–131.

Buhusi CV, Meck WH (2005) What makes us tick? Functional and neural mechanisms of interval timing. Nature Reviews Neuroscience 6:755–765.

Capilla A, Schoffelen J-M, Paterson G, Thut G, Gross J (2014) Dissociated α-Band Modulations in the Dorsal and Ventral Visual Pathways in Visuospatial Attention and Perception. Cerebral Cortex 24:550–561.

Chauvin JJ, Gillebert CR, Rohenkohl G, Humphreys GW, Nobre AC (2016) Temporal orienting of attention can be preserved in normal aging. Psychology and Aging 31:442–455.

Dahl MJ, Ilg L, Li S-C, Passow S, Werkle-Bergner M (2019) Diminished pre-stimulus alpha-lateralization suggests compromised self-initiated attentional control of auditory processing in old age. NeuroImage 197:414–424.

Deiber M-P, Ibañez V, Missonnier P, Rodriguez C, Giannakopoulos P (2013) Age-associated modulations of cerebral oscillatory patterns related to attention control. NeuroImage 82:531–546.

Deng Y, Choi I, Shinn-Cunningham B (2020) Topographic specificity of alpha power during auditory spatial attention. NeuroImage 207:116360.

Dew ITZ, Buchler N, Dobbins IG, Cabeza R (2011) Where Is ELSA? The Early to Late Shift in Aging. Cerebral Cortex 22:2542–2553.

Dimitrijevic A, Smith ML, Kadis DS, Moore DR (2017) Cortical Alpha Oscillations Predict Speech Intelligibility. Frontiers in Human Neuroscience 11:Article 88.

Droit-Volet S, Lorandia F, Coull JT (2019) Explicit and implicit timing in aging. Acta Psychologica 193:180–189.

Fiedler L, Ala TS, Graversen C, Alickovic E, Lunner T, Wendt D (2021) Hearing Aid Noise Reduction Lowers the Sustained Listening Effort During Continuous Speech in Noise—A Combined Pupillometry and EEG Study. Ear & Hearing 42:1590–1601.

Fischl B, Sereno MI, Dale AM (1999a) Cortical surface-based analysis II: inflation, flattening, and a surface-based coordinate system. NeuroImage 9:195–207.

Fischl B, Sereno MI, Tootell RBH, Dale AM (1999b) High-resolution intersubject averaging and a coordinate system for the cortical surface. Human Brain Mapping 8:272–284.

Fitzgibbons PJ, Gordon-Salant S (2010) Behavioral Studies With Aging Humans: Hearing Sensitivity and Psychoacoustics. In: The Aging Auditory System (Gordon-Salant S, Frisina RD, Popper AN, Fay RR, eds), pp 111–134. New York, USA: Springer-Verlag.

Foxe JJ, Simpson GV, Ahlfors SP (1998) Parieto-occipital ∼10Hz activity reflects anticipatory state of visual attention mechanisms. NeuroReport 9:3929–3933.

Fu K-MG, Foxe JJ, Murray MM, Higgins BA, Javitt DC, Schroeder CE (2001) Attention-dependent suppression of distracter visual input can be cross-modally cued as indexed by anticipatory parieto–occipital alpha-band oscillations. Cognitive Brain Research 12:145–152.

Garcés P, López-Sanz D, Maestú F, Pereda E (2017) Choice of Magnetometers and Gradiometers after Signal Space Separation. Sensors 17:2926.

Gatehouse S, Noble W (2004) The speech, spatial and qualities of hearing scale (SSQ). International Journal of Audiology 43:85–99.

Getzmann S, Klatt L-I, Schneider D, Begau A, Wascher E (2020) EEG correlates of spatial shifts of attention in a dynamic multi-talker speech perception scenario in younger and older adults. Hearing Research 398:108077.

Gibbon J (1977) Scalar expectancy theory and Weber’s law in animal timing. Psychological Review 84:279–325.

Glasser MF, Coalson TS, Robinson EC, Hacker CD, Harwell J, Yacoub E, Ugurbil K, Andersson J, Beckmann CF, Jenkinson M, Smith SM, Van Essen DC (2016) A multi-modal parcellation of human cerebral cortex. Nature 536:171–178.

Grondin S (2014) About the (non)scalar property for time perception. Advances in experimental medicine and biology 829:17–32.

Gross J, Kujala J, Hämäläinen MS, Timmermann L, Schnitzler A, Salmelin R (2001) Dynamic imaging of coherent sources: Studying neural interactions in the human brain. Proceedings of the National Academy of Sciences 98:694–699.

Hämäläinen MS, Hari R (2002) Magnetoencephalographic (MEG) Characterization of Dynamic Brain Activation: Basic Principles and Methods of Data Collection and Source Analysis. In: Brain Mapping: The Methods (Toga AW, Mazziotta JC, eds), pp 227–253: Academic Press.

Hämäläinen MS, Hari R, Ilmoniemi RJ, Knuutila J, Lounasmaa OV (1993) Magnetoencephalography – theory, instrumentation, and applications to noninvasive studies of the working human brain. Reviews of Modern Physics 65:413–497.

Han X, Jovicich J, Salat DH, van der Kouwe A, Quinn B, Czanner S, Busa E, Pacheco J, Albert M, Killiany R, Maguire P, Rosas D, Makris N, Dale AM, Dickerson B, Fischl BR (2006) Reliability of MRI-derived measurements of human cerebral cortical thickness: The effects of field strength, scanner upgrade and manufacturer. NeuroImage 32:180–194.

Hari M, PhD, Riitta, Puce P, Aina (2017) MEG-EEG Primer: Oxford University Press.

Heideman SG, Rohenkohl G, Chauvin JJ, Palmer CE, van Ede F, Nobre AC (2018) Anticipatory neural dynamics of spatial-temporal orienting of attention in younger and older adults. NeuroImage 178:46–56.

Henry MJ, Obleser J (2012) Frequency modulation entrains slow neural oscillations and optimizes human listening behavior. Proceedings of the National Academy of Sciences 109:20095–20100.

Henry MJ, Herrmann B, Obleser J (2014) Entrained neural oscillations in multiple frequency bands co-modulate behavior. Proceedings of the National Academy of Sciences 111:14935–14940.

Henry MJ, Herrmann B, Obleser J (2016) Neural microstates govern perception of auditory input without rhythmic structure. The Journal of Neuroscience 36:860–871.

Henry MJ, Herrmann B, Kunke D, Obleser J (2017) Aging affects the balance of neural entrainment and top-down neural modulation in the listening brain. Nature Communications 8:15801.

Herrmann B, Johnsrude IS (2018) Attentional State Modulates the Effect of an Irrelevant Stimulus Dimension on Perception. Journal of Experimental Psychology: Human Perception and Performance 44:89–105.

Herrmann B, Johnsrude IS (2020) A Model of Listening Engagement (MoLE). Hearing Research.

Herrmann B, Maess B, Johnsrude IS (2018) Aging Affects Adaptation to Sound-Level Statistics in Human Auditory Cortex. The Journal of Neuroscience 38:1989–1999.

Herrmann B, Maess B, Johnsrude IS (2022a) Sustained responses and neural synchronization to amplitude and frequency modulation in sound change with age. BioRxiv.

Herrmann B, Maess B, Johnsrude IS (2022b) A Neural Signature of Regularity in Sound is Reduced in Older Adults. Neurobiology of Aging 109:1–10.

Hessler JB, Schäufele M, Hendlmeier I, Nora Junge M, Leonhardt S, Weber J, Bickel H (2017) The 6-Item Cognitive Impairment Test as a bedside screening for dementia in general hospital patients: results of the General Hospital Study (GHoSt). International Journal of Geriatric Psychiatry 32:726–733.

Hong X, Sun J, Bengson JJ, Mangun GR, Tong S (2015) Normal aging selectively diminishes alpha lateralization in visual spatial attention. NeuroImage 106:353–363.

Hornsby BW, Naylor G, Bess FH (2016) A Taxonomy of Fatigue Concepts and Their Relation to Hearing Loss. Ear & Hearing 37:136S–144S.

Humes LE, Busey TA, Craig JC, Kewley-Port D (2009) The effects of age on sensory thresholds and temporal gap detection in hearing, vision, and touch. Attention, Perception, & Psychophysics 71:860–871.

Jensen O, Mazaheri A (2010) Shaping functional architecture by oscillatory alpha activity: gating by inhibition. Frontiers in Human Neuroscience 4:Article 186.

Johnsrude IS, Rodd JM (2016) Factors That Increase Processing Demands When Listening to Speech. In: Neurobiology of Language (Hickok G, Small SL, eds), pp 491–502: Elsevier Academic Press.

Kanwisher N, Wojciulik E (2000) Visual attention: Insights from brain imaging. Nature Reviews Neuroscience 1:91–100.

Klimesch W (1999) EEG alpha and theta oscillations reflect cognitive and memory performance: a review and analysis. Brain Research Reviews 29:169–195.

Klimesch W, Sauseng P, Hanslmayr S (2007) EEG alpha oscillations: The inhibition–timing hypothesis. Brain Research Reviews 53:63–88.

Leenders MP, Lozano-Soldevilla D, Roberts MJ, Jensen O, De Weerd P (2018) Diminished Alpha Lateralization During Working Memory but Not During Attentional Cueing in Older Adults. Cerebral Cortex 28:21–32.

Lehtelä L, Salmelin R, Hari R (1997) Evidence for reactive magnetic 10-Hz rhythm in the human auditory cortex. Neuroscience Letters 222:111–114.

Makeig S, Bell AJ, Jung T-P, Sejnowski TJ (1996) Independent component analysis of electroencephalographic data. In: Advances in Neural Information Processing Systems (Touretzky D, Mozer M, Hasselmo M, eds). Cambridge, MA, USA: MIT Press.

Mattys SL, Davis MH, Bradlow AR, Scott SK (2012) Speech recognition in adverse conditions: A review. Language and Cognitive Processes 27:953–978.

Mok RM, Myers NE, Wallis G, Nobre AC (2016) Behavioral and Neural Markers of Flexible Attention over Working Memory in Aging. Cerebral Cortex 26:1831–1842.

Müller N, Leske S, Hartmann T, Szebényi S, Weisz N (2015) Listen to yourself: The medial prefrontal cortex modulates auditory alpha power during speech preparation. Cerebral Cortex 25:4029–4037.

Niedermeyer E (1990) Alpha-like rhythmical activity of the temporal lobe. Clinical Electrocencephalography 21:210–224.

Niemi P, Näätänen R (1981) Foreperiod and simple reaction time. Psychological Bulletin 89:133–162.

Nobre AC, van Ede F (2018) Anticipated moments: temporal structure in attention. Nature Reviews Neuroscience 19:34–48.

Nobre AC, Correa A, Coull JT (2007) The hazards of time. Current Opinion in Neurobiology 17:465–470.

Nolte G (2003) The magnetic lead field theorem in the quasi-static approximation and its use for magnetoencephalography forward calculation in realistic volume conductors. Physics in Medicine and Biology 48:3637–3652.

Obleser J, Wöstmann M, Hellbernd N, Wilsch A, Maess B (2012) Adverse Listening Conditions and Memory Load Drive a Common Alpha Oscillatory Network. The Journal of Neuroscience 32:12376–12383.

Oostenveld R, Fries P, Maris E, Schoffelen JM (2011) FieldTrip: Open source software for advanced analysis of MEG, EEG, and invasive electrophysiological data. Computational Intelligence and Neuroscience 2011:Article ID 156869.

Palva S, Palva JM (2011) Functional roles of alpha-band phase synchronization in local and large-scale cortical networks. Frontiers in Psychology 2:Article 204.

Payne L, Guillory S, Sekuler R (2013) Attention-modulated Alpha-band Oscillations Protect against Intrusion of Irrelevant Information. Journal of Cognitive Neuroscience 25:1463–1476.

Peelle JE (2018) Listening Effort: How the Cognitive Consequences of Acoustic Challenge Are Reflected in Brain and Behavior. Ear & Hearing 39:204–214.

Pessoa L, Kastner S, Ungerleider LG (2003) Neuroimaging Studies of Attention: From Modulation of Sensory Processing to Top-Down Control. The Journal of Neuroscience 23:3990–3998.

Pichora-Fuller MK (2003) Processing speed and timing in aging adults: psychoacoustics, speech perception, and comprehension. International Journal of Audiology 42:S59–S67.

Pichora-Fuller MK (2016) How Social Psychological Factors May Modulate Auditory and Cognitive Functioning During Listening. Ear & Hearing 37 Suppl 1:92S–100S.

Pichora-Fuller MK, Kramer SE, Eckert MA, Edwards B, Hornsby BWY, Humes LE, Lemke U, Lunner T, Matthen M, Mackersie CL, Naylor G, Phillips NA, Richter M, Rudner M, Sommers MS, Tremblay KL, Wingfield A (2016) Hearing Impairment and Cognitive Energy: The Framework for Understanding Effortful Listening (FUEL). Ear & Hearing 37 Suppl 1:5S–27S.

Power JD, Petersen SE (2013) Control-related systems in the human brain. Current Opinion in Neurobiology 23:223–228.

Richter M, Gendolla GHE, Wright RA (2016) Three Decades of Research on Motivational Intensity Theory: What We Have Learned About Effort and What We Still Don’t Know. In: Advances in Motivation Science (Elliot AJ, ed), pp 149–186. Cambridge, MA, USA: Academic Press.

Rogers CS, Payne L, Maharjan S, Wingfield A, Sekuler R (2018) Older Adults Show Impaired Modulation of Attentional Alpha Oscillations: Evidence From Dichotic Listening. Psychology and Aging 33:246–258.

Rohenkohl G, Nobre AC (2011) Alpha Oscillations Related to Anticipatory Attention Follow Temporal Expectations. The Journal of Neuroscience 31:14076–14084.

Sander MC, Werkle-Bergner M, Lindenberger U (2012) Amplitude modulations and inter-trial phase stability of alpha-oscillations differentially reflect working memory constraints across the lifespan. NeuroImage 59:646–654.

Shenhav A, Musslick S, Lieder F, Kool W, Griffiths TL, Cohen JD, Botvinick MM (2017) Toward a Rational and Mechanistic Account of Mental Effort. Annual Review of Neuroscience 40:99–124.

Snell KB (1997) Age-related changes in temporal gap detection. The Journal of the Acoustical Society of America 101:2214–2220.

Speer NK, Jacoby LL, Braver TS (2003) Strategy-dependent changes in memory: Effects on behavior and brain activity. Cognitive, Affective, & Behavioral Neuroscience 3:155–167.

Strauss DJ, Francis AL (2017) Toward a taxonomic model of attention in effortful listening. Cognitive, Affective & Behavioral Neuroscience 17:809–825.

Strouse A, Ashmead DH, Ohde RN, Grantham DW (1998) Temporal processing in the aging auditory system. The Journal of the Acoustical Society of America 104:2385–2399.

Taulu S, Kajola M, Simola J (2004) Suppression of Interference and Artifacts by the Signal Space Separation Method. Brain Topography 16:269–275.

Taulu S, Simola J, Kajola M (2005) Applications of the Signal Space Separation Method. IEEE Transactions On Signal Processing 53:3359–3372.

Tiihonen J, Hari R, Kajola M, Karhu J, Ahlfors SP, Tissari S (1991) Magnetoencephalographic 10-Hz rhythm from the human auditory cortex. Neuroscience Letters 129:303–305.

Tuladhar AM, Huurne Nt, Schoffelen J-M, Maris E, Oostenveld R, Jensen O (2007) Parieto-Occipital Sources Account for the Increase in Alpha Activity with Working Memory Load. Human Brain Mapping 28:785–792.

Tune S, Wöstmann M, Obleser J (2018) Probing the limits of alpha power lateralisation as a neural marker of selective attention in middle-aged and older listeners. European Journal of Neuroscience 48:2537–2550.

Tune S, Alavash M, Fiedler L, Obleser J (2021) Neural attentional-filter mechanisms of listening success in middle-aged and older individuals. Nature Communications 12:4533.

Upadhyaya AK, Rajagopal M, Gale TM (2010) The Six Item Cognitive Impairment Test (6-CIT) as a screening test for dementia: comparison with Mini-Mental State Examination (MMSE) Current Aging Science 3:138–142.

Vaden RJ, Hutcheson NL, McCollum LA, Kentros J, Visscher KM (2012) Older adults, unlike younger adults, do not modulate alpha power to suppress irrelevant information. NeuroImage 63:1127–1133.

Vallesi A, McIntosh AR, Stuss DT (2009) Temporal preparation in aging: A functional MRI study. Neuropsychologia 47:2876–2881.

van Dijk H, Schoffelen J-M, Oostenveld R, Jensen O (2008) Prestimulus Oscillatory Activity in the Alpha Band Predicts Visual Discrimination Ability. The Journal of Neuroscience 28:1816–1823.

Wearden JH (2003) Applying the scalar timing model to human time psychology: Progress and challenges. In: Time and mind II: Information processing perspectives., pp 21–39. Ashland, OH, US: Hogrefe & Huber Publishers.

Westbrook A, Braver TS (2015) Cognitive effort: A neuroeconomic approach. Cognitive, Affective & Behavioral Neuroscience 15:395–415.

Wilsch A, Henry MJ, Herrmann B, Maess B, Obleser J (2015) Alpha Oscillatory Dynamics Index Temporal Expectation Benefits in Working Memory. Cerebral Cortex 25:1938–1946.

Wöstmann M, Alavash M, Obleser J (2019) Alpha Oscillations in the Human Brain Implement Distractor Suppression Independent of Target Selection. The Journal of Neuroscience 39:9797–9805.

Wöstmann M, Schmitt L-M, Obleser J (2020) Does Closing the Eyes Enhance Auditory Attention? Eye Closure Increases Attentional Alpha-Power Modulation but Not Listening Performance. Journal of Cognitive Neuroscience 32:212–225.

Wöstmann M, Maess B, Obleser J (2021) Orienting auditory attention in time: Lateralized alpha power reflects spatio-temporal filtering. NeuroImage 228:117711.

Wöstmann M, Herrmann B, Wilsch A, Obleser J (2015) Neural Alpha Dynamics in Younger and Older Listeners Reflect Acoustic Challenges and Predictive Benefits. The Journal of Neuroscience 35:1458–1467.

Wöstmann M, Herrmann B, Maess B, Obleser J (2016) Spatiotemporal dynamics of auditory attention synchronize with speech. Proceedings of the National Academy of Sciences 113:3873–3878.

Zanto TP, Gazzaley A (2017) Cognitive Control and the Ageing Brain. In: The Wiley Handbook of Cognitive Control (Egner T, ed), pp 476–490: John Wiley & Sons.

Zanto TP, Pan P, Liu H, Bollinger J, Nobre AC, Gazzaley A (2011) Age-related changes in orienting attention in time. The Journal of Neuroscience 31:12461–12470.

