## Supplemental Materials for "Neural signatures of task-related fluctuations in auditory attention change with age"

Baycrest, Toronto, ON, Canada

[3]Department of Psychology,

University of Toronto, Toronto, ON, Canada

[4]Brain Networks Group,

Max Planck Institute for Human Cognitive and Brain Sciences, Leipzig, Germany

[5]Max Planck Research Group "Neural and Environmental Rhythms",

Max Planck Institute for Empirical Aesthetics, Frankfurt am Main, Germany

[6]Department of Psychology

University of Lübeck, Lübeck, Germany

[7]School of Communication Sciences & Disorders,

The University of Western Ontario, N6A 5B7, London, ON, Canada

*Correspondence concerning this article should be addressed to Björn Herrmann, Rotman Research Institute, Baycrest, 3560 Bathurst St, North York, Ontario, M6A 2E1, Canada.

### Supplementary Results

#### Listening-related alpha-power frequency is smaller than 10 Hz

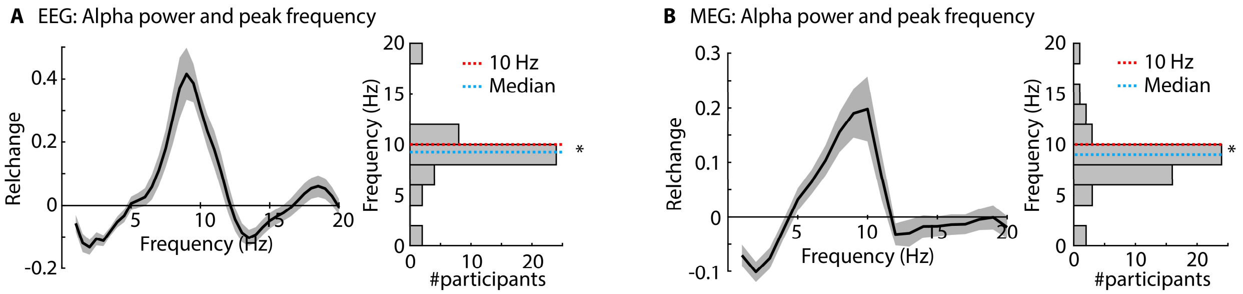


**Figure S1: Alpha power and peak frequency.** Data from the two EEG experiments (A) and one MEG experiment (B) are shown. **Left:** Mean power across participants for different neural frequencies (average of data from -0.8 to -0.2 s prior to target onset). Gray shading reflects the standard error of the mean. **Right:** Histogram of peak frequencies. The peak frequency for both EEG and MEG data was significantly smaller than 10 Hz (EEG: sign-test, $p = 0.0034$; MEG: sign-test, $p = 7 \times 10^{-5}$). $^*p < 0.05$

#### Age differences in alpha power in superior parietal cortex

Analysis of overall age-group differences in alpha power focused on pre-gap alpha power (across cue conditions) and the two regions of interest (superior temporal cortex and superior parietal cortex). Alpha power was larger in superior parietal cortex for younger compared to older adults ($t_{49} = 2.943$, $p = 0.005$, $d = 0.824$), whereas no age difference was found in superior temporal cortex ($t_{49} = 0.741$, $p = 0.462$, $d = 0.208$; Figure S2; Region × Age Group interaction: $F_{1,49} = 8.156$, $p = 0.006$, $\eta_p^2 = 0.143$).

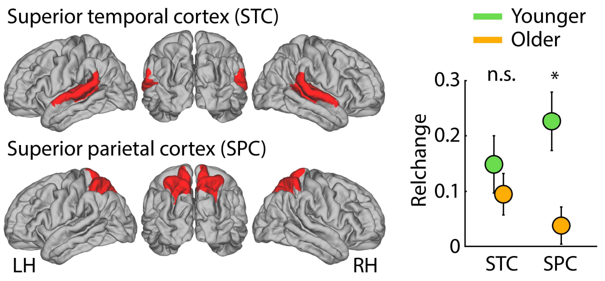


**Figure S2: Age-group differences in alpha power.** Mean pre-gap alpha power (-0.8 to -0.2 s) for two regions of interest: superior temporal cortex (STC; including auditory cortex) and superior parietal cortex (SPC). $^*p \leq 0.05$, n.s. – not significant

### EEG/MEG results for alpha power in the 8–12 Hz frequency window

#### Experiment 1

Alpha power was sensitive to whether participants attended to the first versus second half of the sound: alpha power increased for longer when participants knew the gap would occur later, that is, in the second compared to the first half. For the rmANOVA, we observed a Time Interval × Cue interaction ($F_{1,21}$ = 11.521, p = 0.003, $\eta_p^2$ = 0.354): Alpha power was larger in the first interval (1–1.8 s) compared to the second interval (5.5–6.3 s) when participants were cued to the first half ($F_{1,21}$ = 7.572, p = 0.012, $\eta_p^2$ = 0.265), but alpha power was larger in the second interval (5.5–6.3 s) compared to the first interval (1–1.8 s) when participants were cued to the second half of the sound ($F_{1,21}$ = 8.184, p = 0.009, $\eta_p^2$ = 0.280).

#### Experiment 2

The rmANOVA revealed a Time Interval × Cue interaction ($F_{1,19}$ = 23.385, p = $1.1 \cdot 10^{-4}$, $\eta_p^2$ = 0.552). Alpha power was larger in the first interval (1–1.8 s) compared to the second interval (5.5–6.3 s) when participants were cued to the first half ($F_{1,19}$ = 9.137, p = 0.007, $\eta_p^2$ = 0.325), but alpha power was larger in the second interval (5.5–6.3 s) compared to the first interval (1–1.8 s) when participants were cued to the second half ($F_{1,19}$ = 15.049, p = 0.001, $\eta_p^2$ = 0.442). These results replicate the results from Experiment 1 by showing that alpha power indicates when in time individuals listen attentively.

In addition, the Time Interval × Cue × Task Difficulty interaction was marginally significant ($F_{1,19}$ = 3.734, p = 0.068, $\eta_p^2$ = 0.164): we conducted an exploratory analysis to understand this marginal interaction in more detail. Separately for each cue condition (first half; second half), a rmANOVA with Time Interval (first interval: 1–1.8 s; second interval: 5.5–6.3 s) and Task Difficulty (easy; difficult) factors was conducted. When participants were cued to the first half of the sound, alpha power was larger for difficult compared to easy trials (effect of Task Difficulty: $F_{1,19}$ = 5.826, p = 0.026, $\eta_p^2$ = 0.235), but there was no Time Interval × Task Difficulty interaction (p = 0.334). In contrast, when participants were cued to the second half of the sound, alpha power increased over time in both difficulty conditions, but the increase was steeper for difficult compared to easy trials (Time Interval × Task Difficulty interaction: $F_{1,19}$ = 5.643, p = 0.028, $\eta_p^2$ = 0.229).

#### Experiment 3

##### *Younger and older adults show attention-related alpha power modulations*

MEG data were analyzed in sensor space to examine whether MEG alpha power is sensitive to when participants attend. As for Experiments 1 and 2, the Time Interval × Cue interaction ($F_{1,50} = 12.214$, $p = 0.001$, $\eta_p^2 = 0.196$) showed that alpha power was larger in the first interval (1-1.8 s) compared to the second interval (5.5-6.3 s) when participants were cued to the first half ($F_{1,51} = 10.475$, $p = 0.002$, $\eta_p^2 = 0.170$), whereas alpha power was larger in the second compared to the first interval when participants were cued to the second half ($F_{1,51} = 9.050$, $p = 0.004$, $\eta_p^2 = 0.151$). This interaction indicates that alpha power was sensitive to whether participants attended to the first versus second half of the sound.

We also observed a Cue × Age Group interaction ($F_{1,50} = 5.019$, $p = 0.030$, $\eta_p^2 = 0.091$): Averaged across the two analysis time intervals (first and second interval), alpha power was numerically larger for younger ($p = 0.092$) but smaller for older adults ($p = 0.168$) when they were cued to the first compared to the second half of the sound. None of the other effects were significant.

##### *Age-differences in spatial distribution of alpha activity*

The analysis of age-group differences in the spatial configuration of alpha power revealed that peak alpha power was consistently more ventro-lateral in older compared to younger adults (x-axis of left hemisphere: $t_{49} = 2.761$, $p = 0.008$, $d = 0.773$; x-axis of right hemisphere: $t_{49} = 1.04$, $p = 0.303$, $d = 0.291$; z-axis of left hemisphere: $t_{49} = 3.535$, $p = 9 \cdot 10^{-4}$, $d = 0.99$; z-axis of right hemisphere: $t_{49} = 3.942$, $p = 2.6 \cdot 10^{-4}$, $d = 1.104$).

Correlations of activation maps among participants further highlight age differences in the spatial configuration of brain activations: Correlations of spatial alpha-power maps were larger among members of the same age group (within) compared to members of the other age group (between; younger adults: $t_{24} = 2.642$, $p = 0.014$, $d = 0.528$; older adults: $t_{25} = 5.029$, $p = 3.5 \cdot 10^{-5}$, $d = 0.986$). These results suggest that the source configuration underlying listening-related alpha activity differs between younger and older adults.

*Attentional modulation of alpha power in superior parietal cortex is reduced in older adults*

To statistically quantify age-group differences in attentional modulation of alpha activity, we conducted an rmANOVA with factors Cue (first half; second half), Time Interval (first interval: 1–1.8 s; second interval: 5.5–6.3 s), Region (STC; SPC), and Age Group (younger; older). The Cue × Time Interval × Region × Age Group interaction was significant ($F_{1,49} = 14.282$, $p = 4.3 \times 10^{-4}$, $n_p^2 = 0.226$). In younger adults, the Cue × Time Interval × Region interaction was significant ($F_{1,24} = 13.621$, $p = 0.001$, $n_p^2 = 0.362$), showing that alpha power in SPC (Cue × Time Interval: $F_{1,24} = 13.893$, $p = 0.001$, $n_p^2 = 0.367$), and only to a lesser degree STC (Cue × Time Interval: $F_{1,24} = 6.032$, $p = 0.022$, $n_p^2 = 0.201$), decreased from the 1–1.8 s (first interval) to 5.5–6.3 s (second interval) when participants were cued to the first half of the sound, and increased from the 1–1.8 s to 5.5–6.3 s intervals when participants were cued to second half of the sound. In older adults, the Cue × Time Interval interaction ($F_{1,25} = 12.451$, $p = 0.002$, $n_p^2 = 0.332$) was not further specified by brain region (Cue × Time Interval × Region: $F_{1,25} = 2.901$, $p = 0.101$, $n_p^2 = 0.104$).

To examine the Cue × Time Interval × Region × Age Group interaction in a different way, we also calculated an attentional modulation index (AMI) as the sum of the difference between time intervals (i.e., difference between first and second interval for cued to first half, plus the difference between second and first interval for cued to second half), separately for each participant and region. The attentional modulation was larger in SPC for younger than older adults ($t_{49} = 2.008$, $p = 0.050$, $d = 0.562$), whereas there was no difference between age groups for STC ($t_{49} = -0.849$, $p = 0.400$, $d = 0.238$; Region × Age-Group interaction: $F_{1,49} = 14.282$, $p = 4.3 \times 10^{-4}$, $n_p^2 = 0.226$).

#### Experiment S1: Resting-state alpha activity does not differ between younger and older adults

We have shown in Experiment 3 that listening-related alpha activity relies on temporal cortex and parietal cortex. These regions appear to differ from traditional resting-state alpha activity at around 10 Hz that is largest when individuals have their eyes are closed compared to when their eyes are open (Berger, 1929; van Dijk et al., 2008; Kwok et al., 2019; Wöstmann et al., 2020). Experiment S1 aims to directly test using the same MEG recording setup as for Experiment 3 whether listening-related alpha activity and resting-state alpha activity rely on different brain regions. Moreover, Experiment 3 has revealed differences between younger and older adults in neural circuits underlying listening-related alpha activity. Experiment S1 was also conducted to test whether the age differences are also present for resting-state alpha activity.

#### Methods

##### *Participants*

Resting state MEG data were recorded from 19 younger (age range: 21–31 years; median: 25 years; 11 females and 8 males) and 19 older adults (age range: 54–71 years; median: 63.5 years; 9 females and 10 males) who did not participate in Experiment 3.

##### *Recording and analysis of resting-state data*

Resting-state activity was recorded using the same MEG data-recording settings as described for Experiment 3. Resting-state blocks were 6-min long, during which participants were instructed auditorily via in-ear phones to either close or open their eyes, three times each, for about one minute at the time. Whether participants started with eyes open or closed was counter-balanced across participants.

Artifacts such as blinks, eye-movements, and muscle activity were removed using independent components analysis (Oostenveld et al., 2011). 100 artifact-free data snippets of 7-s duration were randomly selected from each one-minute eyes open/closed recording period (Kwok et al., 2019). A fast Fourier transform (FFT) was calculated for each 7-s snippet and channel (Hann taper, zero-padding). The power spectrum was calculated as the squared magnitude of the resulting complex numbers. Power spectra were averaged across data snippets, independently for eyes-open and eyes-closed conditions. In order to analyze alpha activity, power in the 8–12 Hz frequency band was averaged for each condition. An ANOVA with the within-participants factor Condition (eyes open; eyes closed) and the between-participants factor Age Group (younger; older) was calculated.

In order to compare spatial locations of activity for listening-related and resting-state alpha power, we identified the MEG channel exhibiting the maximum alpha power in MEG data, separately for listening-related alpha power from Experiment 3 and resting-state alpha power from Experiment S1. We tested whether resting-state power originates from more posterior (occipital) sources compared to listening-related power (parietal) by comparing the y-coordinate of the activation maximum using an independent-samples t-test. Age-group differences in the location of peak resting-state alpha activity were assessed using an independent-samples t-test.

*MEG source reconstruction*

Volume conductor and sources models were calculated as for Experiment 3. For source localization of resting-state activity, the cross-spectral density matrix was calculated from the complex coefficients from the fast Fourier transform in the 8–12 Hz frequency band, and source reconstructions were calculated using the DICS beamformer (Gross et al., 2001; real-numbered spatial filters using the dominant direction). Complex coefficients were projected through the filter and power was calculated as the squared magnitude of the coefficients in source space. No MRI scan was available for three younger and eight older adults who participated in the resting sessions, and data from these persons were thus not considered for source analyses (resulting in source-localization datasets from N=16 younger and N=11 older participants).

Results and discussion

As expected, alpha power (8–12 Hz) was larger when participants had their eyes closed compared to open ($F_{1,36}$ = 22.693, p = 3.1 × $10^{-5}$, $\eta_p^2$ = 0.387; Figure S3A; Berger, 1929; van Dijk et al., 2008; Kwok et al., 2019; Wöstmann et al., 2020). Alpha power was smaller in older compared to younger adults ($F_{1,36}$ = 6.816, p = 0.013, $\eta_p^2$ = 0.159; Figure S3A), but there was no Age Group × Condition interaction ($F_{1,36}$ = 0.639, p = 0.429, $\eta_p^2$ = 0.017).

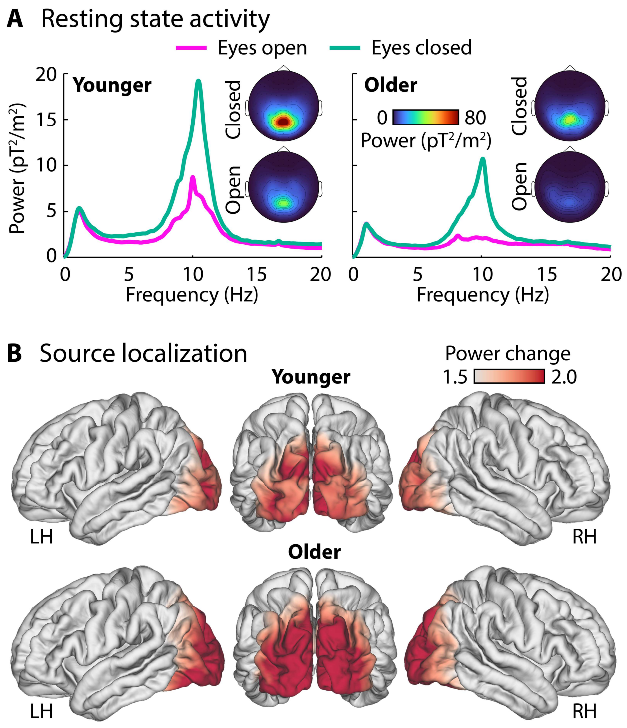


**Figure S3: Resting state activity. A:** MEG power spectra under resting: eyes closed and eyes open conditions for younger (N=19) and older individuals (N=19). **B:** Source localization (ratio of eyes-closed to eyes-open) for resting-state alpha activity (8–12 Hz) for each age group. Source localizations are based on a subset of 16 younger and 11 older participants for who MRI scans were available.

Topographical distributions as well as source localizations show that occipital cortex underlies resting-state alpha activity (Figure S3B). This is different from the parietal and temporal cortices that we identified as generating listening-related alpha activity (Figure 6). Indeed, the peak activation maximum of alpha power was more posterior for resting-state alpha activity compared to listening-related alpha activity ($t_{88}$ = 8.926, p = 5.83 × $10^{-14}$, d = 1.905; sensor-space data were used for this analysis, because brain anatomical information was available only for a limited number of individuals participating in the resting-state experiment). The spatial location of peak resting-state alpha activity did not differ between age groups ($p > 0.15$).

Consistent with the suggestion of multiple alpha oscillators in the brain (Başar et al., 1997; Klimesch, 1999; Bollimunta et al., 2008; van Dijk et al., 2008; Wöstmann et al., 2020), our data suggest distinct neural alpha generators underlying resting-state and listening-related alpha activity and that differences in activated brain regions between age groups are specific to listening-related alpha and do not generalize to resting-state alpha activity.

### References

Başar E, Schürmann M, Başar-Eroglu C, Karakaş S (1997) Alpha oscillations in brain functioning: an integrative theory. International Journal of Psychophysiology 26:5-29.

Berger H (1929) Über das Elektrenkephalogramm des Menschen. European Archives of Psychiatry and Clinical Neuroscience 87:527-570.

Bollimunta A, Cheng Y, Schroeder CE, Ding M (2008) Neuronal Mechanisms of Cortical Alpha Oscillations in Awake-Behaving Macaques. The Journal of Neuroscience 28:9976-9988.

Gross J, Kujala J, Hämäläinen MS, Timmermann L, Schnitzler A, Salmelin R (2001) Dynamic imaging of coherent sources: Studying neural interactions in the human brain. Proceedings of the National Academy of Sciences 98:694-699.

Klimesch W (1999) EEG alpha and theta oscillations reflect cognitive and memory performance: a review and analysis. Brain Research Reviews 29:169-195.

Kwok EYL, Cardy JO, Allman BL, Allen P, Herrmann B (2019) Dynamics of spontaneous alpha activity correlate with language ability in young children. Behavioural Brain Research 359:56-65.

Oostenveld R, Fries P, Maris E, Schoffelen JM (2011) FieldTrip: Open source software for advanced analysis of MEG, EEG, and invasive electrophysiological data. Computational Intelligence and Neuroscience 2011:Article ID 156869.

van Dijk H, Schoffelen J-M, Oostenveld R, Jensen O (2008) Prestimulus Oscillatory Activity in the Alpha Band Predicts Visual Discrimination Ability. The Journal of Neuroscience 28:1816-1823.

Wöstmann M, Schmitt L-M, Obleser J (2020) Does Closing the Eyes Enhance Auditory Attention? Eye Closure Increases Attentional Alpha-Power Modulation but Not Listening Performance. Journal of Cognitive Neuroscience 32:212-225.
